# Intermittent Fasting induced ketogenesis inhibits mouse epithelial ovarian tumors by promoting anti-tumor T cell response

**DOI:** 10.1101/2023.03.08.531740

**Authors:** Mary Priyanka Udumula, Harshit Singh, Rashid Faraz, Laila Poisson, Nivedita Tiwari, Irina Dimitrova, Miriana Hijaz, Radhika Gogoi, Margaret Swenor, Adnan Munkarah, Shailendra Giri, Ramandeep Rattan

## Abstract

Epithelial Ovarian Cancer (EOC) is the most lethal gynecologic cancer with limited genetic alterations identified that can be therapeutically targeted. In tumor bearing mice, short-term fasting, fasting mimicking diet and calorie restriction enhance the activity of antineoplastic treatment by modulating systemic metabolism and boosting anti-tumor immunity. We tested the outcome of sixteen-hour intermittent fasting (IF) on mouse EOC progression with focus on fasting driven antitumor immune responses. IF resulted in consistent decrease of tumor promoting metabolic growth factors and cytokines, recapitulating changes that creates a tumor antagonizing environment. Immune profiling revealed that IF profoundly reshapes anti-cancer immunity by inducing increase in CD4^+^ and CD8^+^ cells, paralleled by enhanced antitumor Th1 and cytotoxic responses, by enhancing their metabolic fitness. Metabolic studies revealed that IF generated bioactive metabolite BHB which can be a potential substitute for simulating the antitumor benefits of IF. However, in a direct comparison, IF surpassed exogenous BHB therapy in improving survival and activating anti-tumor immune response. Thus, our data provides strong evidence for IF and its metabolic mediator BHB for ameliorating EOC progression and as a viable approach in maintaining and sustaining an effective anti-tumor T cell response.

## Introduction

Epithelial ovarian cancer (EOC), the most prevalent form of ovarian cancer, is the primary cause of mortality due to gynecologic cancer in the United States. Despite breakthroughs in therapy choices for EOC, its prognosis remains dismal, with a survival rate that has remained unchanged for decades at 45%, and approximately 20% for those diagnosed at a late stage, making it the fifth leading cause of cancer-related death in women^1^. The multimodal treatment for EOC consists of maximal surgical resection followed by chemotherapy with platinum and taxanes. The recent introduction of PARP inhibitors and anti-angiogeneic treatments has provided some advantages. Despite the fact that the majority of patients initially respond to the treatments, more than 80% will reccur within two years, limiting the benefits of treatment. In addition, the financial toxicity associated with chemotherapy is an added burden for EOC patients, diminishing their quality of life (QoL)^2, 3^. Consequently, it is of the utmost importance to identify complementary approaches that can improve therapy efficacy while causing minimal toxicity. An understudied and undervalued area in EOC is the modulation of environmental factors that play a key role in determining the balance between disease and outcome. Among the numerous environmental influences, nutrition has been demonstrated to influence disease development, therapeutic response, and survival in a variety of cancers^4, 5^, but such evidence is scarce in EOC. In addition, there are currently no universally accepted dietary recommendations for EOC patients that might improve their prognosis and survival.

Dietary restriction is one of the oldest diet-based interventions applied and tested in cancer and other disorders^6, 7^. It has been demonstrated that caloric restriction reduces the risk of several metabolic illnesses, enhances life expectancy, and inhibits cancer growth^8, 9^. Recent studies have indicated that the ketogenic diet, which severely restricts carbohydrates and causes ketosis, can improve the prognosis of numerous diseases and cancer^10, 11, 12^. Fasting is another approach to achieve ketosis, with intermittent fasting (IF) being the most popular, extensively used, and scientifically proven to confer similar health benefits, particularly on cancer growth^13^. IF can induce metabolic responses and adaptations that can have far-reaching effects on a patient’s physiology, including the rate of tumor progression, treatment response, anti-tumor immune response, and QoL^7, 13, 14, 15^. While long-term fasting schedules have not been examined in cancer patients, a small number of clinical trials have shown the combination of short-term fasting (24-72h fast) with chemotherapy, to be feasible, safe and result in reduced chemotherapy side effects and better QoL measures^16, 17,18^.

Recent recognition of reprogramming of immunometabolism as a crucial event regulating the function of all immune cells has pushed research into dietary interventions that influence the immune system. Recent cancer studies have eloquently demonstrated the ability of ketogenic and other restrictive diets to stimulate an efficient anti-tumor immune response^11, 19, 20^. None of these investigations, however, have been conducted in EOC. Here, we assessed the effect of a 16-hour fast and 8-hour feeding IF on survival and immune response against the tumor in preclinical models of syngeneic EOC. We found that IF increases overall survival via enhancing T-cell antitumor immunity, and via buildup of the ketone body BHB (ß-hydroxy butyrate), regardless of the tumor’s mutational signature. Our research demonstrates the efficacy of IF as an adjuvant intervention and a strategy to improve antitumor immune response in EOC.

## Methods

### Cell lines and reagents

ID8 ^p53+/+^, TRP53 mutated (ID8 ^p53-/-^) and PTEN mutated (ID8^p53-/-, PTEN-/-^) ID8 cells were a gift from Dr. Ian McNeish (Imperial College, London)^21^. All the cell lines were cultured in Dulbecco’s modified Eagles media (Hyclone, Logan, UT), 5% FBS (BioAbChem, Ladson, SC), 4% Insulin,-Transferrin-Selenium (Invitrogen), 100 U/ml penicillin and 100 U/ml streptomycin (Hyclone, Logan, UT), 2mM L-glutamine, 1mM sodium pyruvate. All cells were cultured at 37°C in a humidified 5% CO2 incubator. DL-β hydroxybutyric (BHB) acid sodium salt was purchased from Sigma-Aldrich (Burlington, MA).

### Animal model and experiments

#### Induction of intra-peritoneal ovarian tumor

C57/B6 female mice of 6-8 weeks were acquired from Jackson Laboratory (Bar Harbor, ME) and were maintained under standard optimized conditions in the animal facility at Henry Ford Hospital (Detroit, MI). Prior to the start of the study, mice were acclimated in house for one week. To induce ovarian cancer, mice were injected intraperitoneally (IP) with 10 x 10^6^ ID8 cells (ID8^p53+/+^ or ID8^p53-/-^ or ID8^p53-/- PTEN-/-^) in 200μl of PBS. After one week of tumor induction mice were randomly divided into regular diet (RD) or intermittent fasting (IF) groups. Monitoring for tumor burden was performed by tracking weight, abdominal circumference, and ascites formation as before^22, 23^. Mice that experienced severe signs of clinical distress (cachexia, anorexia, increased respiration, excess ascites) or when abdominal circumference exceeded 8 cm, or impaired mobility or bodily functions, were immediately euthanized.

#### Induction of subcutaneous ovarian tumor

Subcutaneous ovarian tumors were initiated by injecting 2×10^6^ ID8^p53-/-^ cells suspended in 150μl of 50% Matrigel (Corning Life Science, Massachusetts) in the subcutaneous layer of the left flank of the mice. Once visible, the tumors were measured using a digital Vernier caliper. Tumor volume was calculated by using the standard formula 1/2(L x W)^2^, where L is the longest and W is the shorter length measured^24^. After the formation of visible tumors, tumor volume was assessed weekly.

#### Intermittent Fasting protocol

IF intervention consisted of 16 hours of fasting and 8 hours of *ad libitum* feeding for five consecutive days and two consecutive days of 24-hour *ad libitum* feeding during the week. The regular diet (RD) control group received 24 hour, 7 days a week *ad libitum* feeding throughout the study. Continuous access to water was provided to all mice throughout the study. IF was initiated one week after tumor injections and mice were monitored every other day till the end of the study. Food intake was measured over a period of one week at multiple time points during the study.

#### Treatments

All treatments were started after one week of tumor injection and each treatment group consisted of 10 mice. Immunotherapy groups received *InVivoMab* anti-mouse PD-1 or control *InVivoMab* rat IgG2b isotype given IP once a week at 100 μg/mice for 4 weeks alone or in combination with IF. In depletion experiments, mice were treated with *InVivo*Mab anti-mouse CD8α (200μg/mice) or *InVivoPlus* anti-mouse CD4 (200μg/mice) or *InVivo*Mab rat IgG2b (BioXcell, Lebanon, NH) twice via IP injections per week for a total of 6 doses. Depletion efficacy was measured by flow cytometry after 4 doses of antibody injections. All antibodies (Table S1) were purchased from BioXcell (West Lebanon, NH). BHB treatments were given IP at the dose of 300mg/kg bd wt thrice weekly, while the control mice received PBS.

#### Survival Curve

For estimating overall survival, n=12 mice per group were injected with respective ID8 cells and tumors were allowed to proceed until the abdominal circumference reached 8 cm, according to the Hospital Institutional Animal Care and Use Committee (IACUC) approved end point. The mice were then humanely euthanized. Survival curves were generated using Kaplan-Meir analysis using Prism 8 (GraphPad Software, La Jolla, CA).

#### Ethics statement

All protocols were approved by the Henry Ford Hospital IACUC prior to any experiments. All institutional and national guidelines for the care and use of laboratory animals were followed.

### Fluorescence-activated cell sorting analysis

Surface staining (CD45, CD3, CD4, CD8) and intracellular staining (IFN-γ, IL4, Granzyme B and Perforin) was performed in ascites and blood according to manufacturer’s recommended dilution and as published before^23^. Flow cytometric analysis was performed on a BD FACS Calibur (BD Biosciences, San Jose, CA, USA), and results were analyzed using Flowjo software (Version 10.8.1; BD Biosciences, San Jose, CA, USA). T-Distributed Stochastic Neighbor Embedding (tSNE) clusters were prepared using Flowjo software (Version 10.8.1)^25, 26^. All fluorochrome labeled antibodies (Table S2) were purchased from Biolegend (San Diego, CA, USA). Details are described in supplementary methods.

### ELISA

The levels of IL-6, IL-1β, IL-4, IL-10, IFNγ, MCP-1, TNFα (Biolegend, San Diego, CA, USA), GM-CSF, IGF-1, Leptin, Adiponectin, Insulin (R&D Biosystems Minneapolis, MN, USA), Pre-albumin (Crystal Chem Inc., El Grove Village, IL, USA), and BHB (Cayman Chemical, Ann Arbor, MI, USA) in ascites, plasma or cell culture supernatants were measured according to manufacturers’ instructions as before^23^. ELISA kits are listed in table S3.

### Adipokine array

Ascites collected from RD and IF mice (n=4/ group) were pooled and diluted 1:3 and applied in the adipokine array (ARY-013, R&D systems, Minneapolis, MN, USA). The proteome profiler adipokine array detects 38 adipokines (Table S4) in duplicate on nitrocellulose membranes. Expression of the adipokines were analyzed by Quick spots software designed for proteome arrays (R&D Systems, Minneapolis, MN, USA).

### Realtime PCR

RNA was extracted (Qiagen, Valencia, CA, USA) and quantified (Qubit Fluorometer, Invitrogen, Waltham, MA, USA) from liver, and purified CD4^+^ and CD8^+^ T-cells prior to quantifying mRNA expression as described before^23^ and in supplementary methods. Primer (Integrated DNA Technologies, Coralville, IA) sequences for acetyl coenzyme A acetyltransferase (ACAT), 3-Hydroxy-3-methylglutaryl-CoA synthase (HMGCS2), 3-Hydroxy-3-methylglutaryl-CoA lyase (HMGCL), beta-hydroxybutyrate dehydrogenase 1 (BDH1), Succinyl-CoA:3-ketoacid CoA transferase (SCOT) and G protein-coupled receptor 109a (GPR109a, also known as Hydroxycarboxylic acid receptor 2, HCA_2_) are listed in table S5.

### Seahorse metabolic analysis

CD4^+^ and CD8^+^ cells were isolated from spleens of RD and IF tumor bearing mice using BD-IMAG anti-mouse CD4 and CD8 magnetic particles (BD Biosciences, San Jose, CA) and were plated at a density of 7 x10^5^ cells/well in cell-tak coated XFe 96 cell plates. Oxygen consumption rate (OCR) and Extracellular acidification (ECAR) rate were measured using XFe 96 seahorse analyzer (Agilent, Santa Clara, CA, USA) and analyzed as describe before^23, 27^ and detailed in supplementary methods.

### *Ex-Vivo* co-culture experiments

CD8^+^ and CD4^+^ T-cells isolated from spleens of naïve mice using BD-IMAG anti-mouse CD4^+^ and CD8^+^ magnetic particles (BD Biosciences, San Jose, CA, USA) were cultured in 96-well plates (Nunc, Wiesbaden, Germany) at a density of 3×10^5^ cells/well in 200μL RPMI 1640, supplemented with 5% serum, 100 U/mL penicillin, 100 μg/mL streptomycin, 2 mM L-glutamine, and 1 mM sodium pyruvate. The cells were activated for 72 hours in presence of recombinant mouse IL-2 (10 ng/ml) (BD Pharmingen Biosciences, San Diego, CA, USA) and Gibco Dyna mouse CD3/CD28 beads (Thermo Fischer, Detroit, MI, USA). One set was treated with 10mM BHB, and after 72 hours, 7AAD (eBioscience, San Diego, CA, USA) labeled 3×10^5^ ID8^p53-/-^ tumor cells/well were added to the plates for another 72 hours. Flow cytometry was performed to enumerate 7AAD^+^ tumor cells and the unlabeled T cells^28^. IFNγ release in the cell supernatant was determined by ELISA after 48 hours of co-culture.

### Metabolomics

Metabolomic profiling was performed by Metabolon Inc. (Morrisville, MA, USA). Statistical and bioinformatic analysis was performed by the Bioinformatics core at HFH as previously described^29, 30, 31^.

### Quantitation of BHB by LC-MS/MS

The levels of BHB in ascites and plasma of untreated, BHB treated and IF mice were quantified by ultrahigh performance liquid chromatography-tandem mass spectroscopy (UPLC-MS/MS) (Waters, Milford, MA, USA) as described before^31^ and detailed in supplementary methods.

### Statistical Analysis

An unpaired students t-test or one-way ANOVA was used where appropriate. Kaplan Meier analysis was used to determine the survival curve. The significance of survival curves was estimated by using the Gehan Breslow-Wilcoxon test. All analyses were carried out using Prism 8 (GraphPad Software, La Jolla, CA).

## Results

### IF improves overall survival in EOC mouse models irrespective of mutational markers

We compared the anti-tumor effects of a 16-hour IF intervention to those of a non-fasting regular diet (RD) control group in C57/B6 mice injected intraperitoneally (IP) with syngeneic ID8 ovarian tumor cells (ID8^p53+/+^ or ID8^p53-/-^ or ID8^p53-/-, PTEN-/-^), a widely used transplantable murine model to generate peritoneal ovarian tumors^21, 32^. Survival was significantly improved by 27 days in IF mice with ID8^p53+/+^ tumors (median survival 86.5 days) compared to RD (median survival 69 days) (Fig. 1A). The improved survival was accompanied by a relative stabilized weight trend in the IF mice, as opposed to the RD mice, whose weights increased (Fig. 1B). The IF mice had a smaller abdominal circumference as a surrogate for ascites burden (Fig. 1C) and a smaller volume of accumulated ascites (Fig. 1D) than the RD mice. Mice with ID8^p53-/-^ tumors, indicative of the aggressive HGSOC^21, 33^, exhibited an improvement of 19 days when subjected to IF (median survival 70 days) compared to mice with RD tumors (median survival 51 days) (Fig. 1E). Similarly, the IF mice exhibited a stable weight pattern, decreased abdominal circumference and ascites volume (Fig. 1F-H). Mice with ID8^p53-/-, PTEN-/-^ tumors, which are indicative of enhanced aggressiveness^21^, showed an improvement of 20 days when subjected to IF (median survival 58 days vs 38 days in the RD group) (Fig. 1I). Likewise, the IF mice exhibited a stable weight trend, decreased abdominal circumference, and decreased ascites volume (Fig. 1J-L). To determine the visual impact on tumors, we implanted subcutaneous ID8^p53-/-^ tumors and subjected a subset of mice to IF. In the control group, tumors were measurable beginning on day 35, but tumors in the IF group did not become measurable until about 42 days and grew more slowly as represented by the ultimate tumor size and tumor wet weight at endpoint (Fig. 1M-P). Food intake measurements revealed that mice treated to IF ingested nearly the same quantity of food in the 8-hour feeding window that the control RD animals ingested daily when fed *ad libitum* (Fig. S1A-C). In addition, albumin levels assessed after 4-5 weeks of continuous IF and used as a starvation marker^34^ were comparable between the IF and RD groups (Fig. S1D-F).

**Fig 1:**
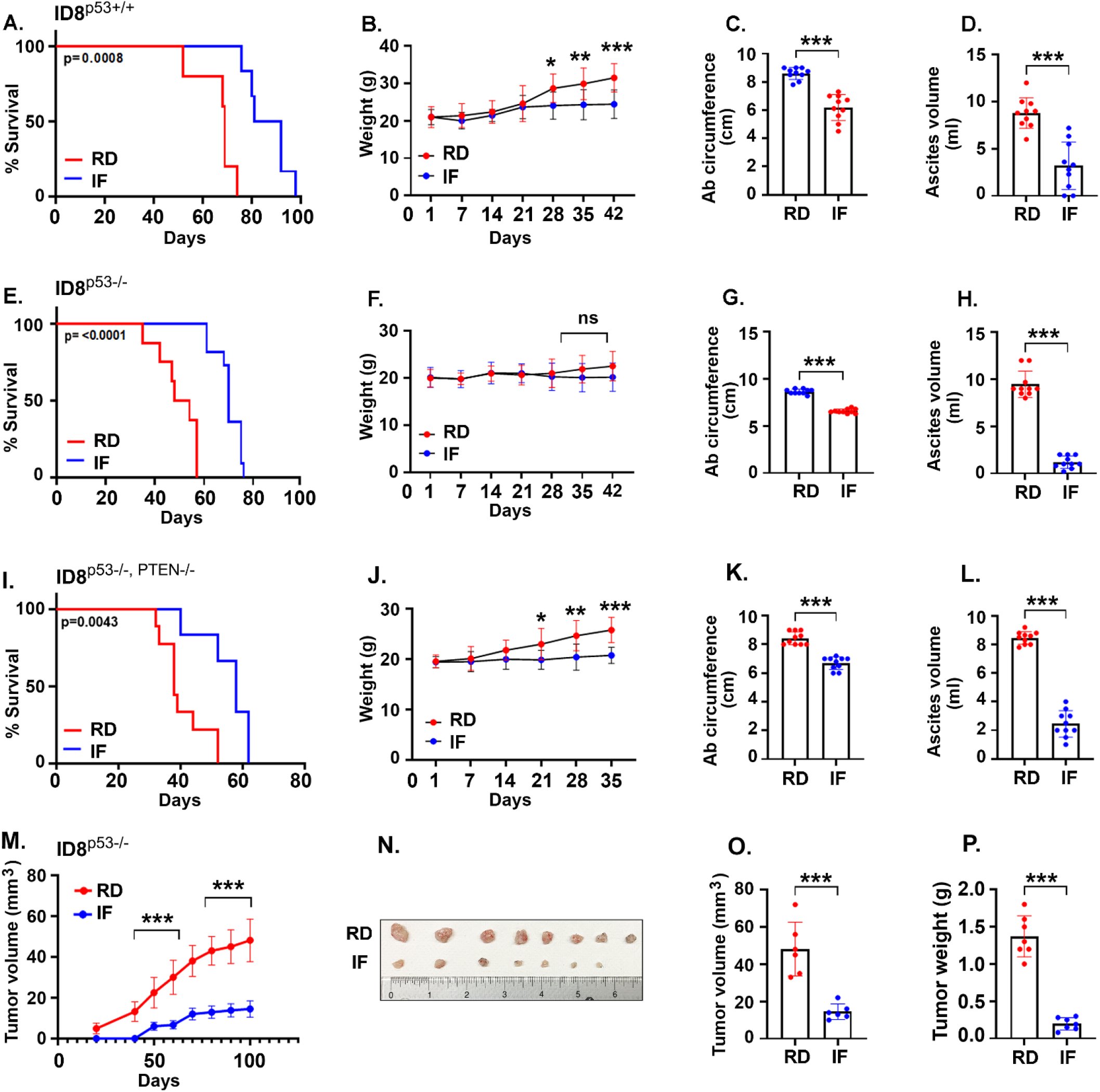
IF improves overall survival in EOC mouse models irrespective of mutational markers. ID8 mouse ovarian cancer cells with and without mutations were injected intraperitoneally into female C57BL/6 mice. After 1 week, mice were randomly subjected to regular ad libitum feeding (RD) or 16-hour intermittent fasting (IF) for 5 days a week and monitored for survival. Kaplan Meier graphs indicating overall survival in mice bearing (A) ID8^p53+/+^ EOC subjected to RD or IF (n=12/group), p=0.0008 by Gehan-Breslow-Wilcoxon test; (E) ID8^p53-/-^ EOC subjected to RD or IF (n=12/group), p<0.0001 by Gehan-Breslow-Wilcox test and (I) ID8^p53-/-,PTEN-/-^ EOC subjected to RD or IF (n=12/group), p=0.0043 by Gehan-Breslow-Wilcox test. Average weekly body weight progression in (B) ID8^p53+/+^, (F) ID8^p53-/-^ and (J) ID8^p53-/-, PTEN-/-^ EOC bearing mice. (C, G, K) Bar graph represents average abdominal circumference at 6 weeks of ID8^p53+/+^, and at 5 weeks of ID8^p53-/-^ and ID8^p53-/-, PTEN-/-^ EOC bearing mice respectively in response to IF. (D, H, L) Ascites volume collected at 6 weeks from ID8^p53+/+^, and at 5 weeks from ID8^p53-/-^ and ID8^p53-/-, PTEN-/-^ EOC bearing mice respectively in response to IF. (M-P) ID8^p53-/-^ cells were injected subcutaneously into the left flank of C57BL/6 mice. After 1 week, mice were subjected to RD and IF and observed for growth of subcutaneous tumors (n=8/group). (M) Line graph represents tumor growth, (N) Photograph of dissected tumors isolated, (O) Bar graph represents tumor volume at the end point of day 100 and, (P) wet tumor weights measured on day 100 from RD and IF mice. ***p < 0.001, ****p<0.0001, IF compared with RD group by Students T-test.

In agreement with previously published findings on other tumor models^35, 36^, we demonstrate for the first time that IF intervention prolongs survival and slows the growth of tumors in syngeneic models of EOC.

### IF decreases tumor promoting growth and inflammatory factors in EOC mouse models

Dietary restrictions have been associated to a reduction in systemic pro-tumor and inflammatory factors^22, 37^. To evaluate the effect of IF on the tumor microenvironment, an adipokine array was performed on the ascites of RD and IF mice with ID8^p53-/-^ tumors (Fig S2). IF decreased tumor-promoting metabolic growth factors such as IGF-1 and leptin, as well as major pro-tumor and inflammatory factors such as FGF acidic, VEGF, ANGPT, MCP-1, M-CSF, and lipocalin, whereas adiponectin levels rose (Fig S2). Validation by ELISA in the ascites and plasma of RD and IF mice, revealed a reduction in growth factors including IGF-1 (Fig 2A, M), insulin (Fig 2B, N) and leptin (Fig 2C, O) and an increase in adiponectin (Fig 2D, P) by IF. Estimation of pro- and anti-inflammatory markers in ascites and plasma revealed that IFNγ, one of the essential cytokines for the cytotoxic CD8^+^ T cells was elevated in the ascites from IF mice compared to RD mice (Fig 2E), but plasma levels remained relatively unaltered (Fig 2Q). Another important anti-inflammatory cytokine IL-10 was also reduced in the ascites and plasma (Fig 2F, U) while other pro-inflammatory cytokines including TNFα, IL-6, IL-4, IL-1β, MCP-1 and GMCSF were significantly lowered in the ascites and plasma of IF mice (Fig 2G-L, 2S-X). Similar differential regulation by IF was also observed in the ascites and plasma of mice with ID8^p53-/-, PTEN-/-^ tumors (Fig S3).

**Fig 2:**
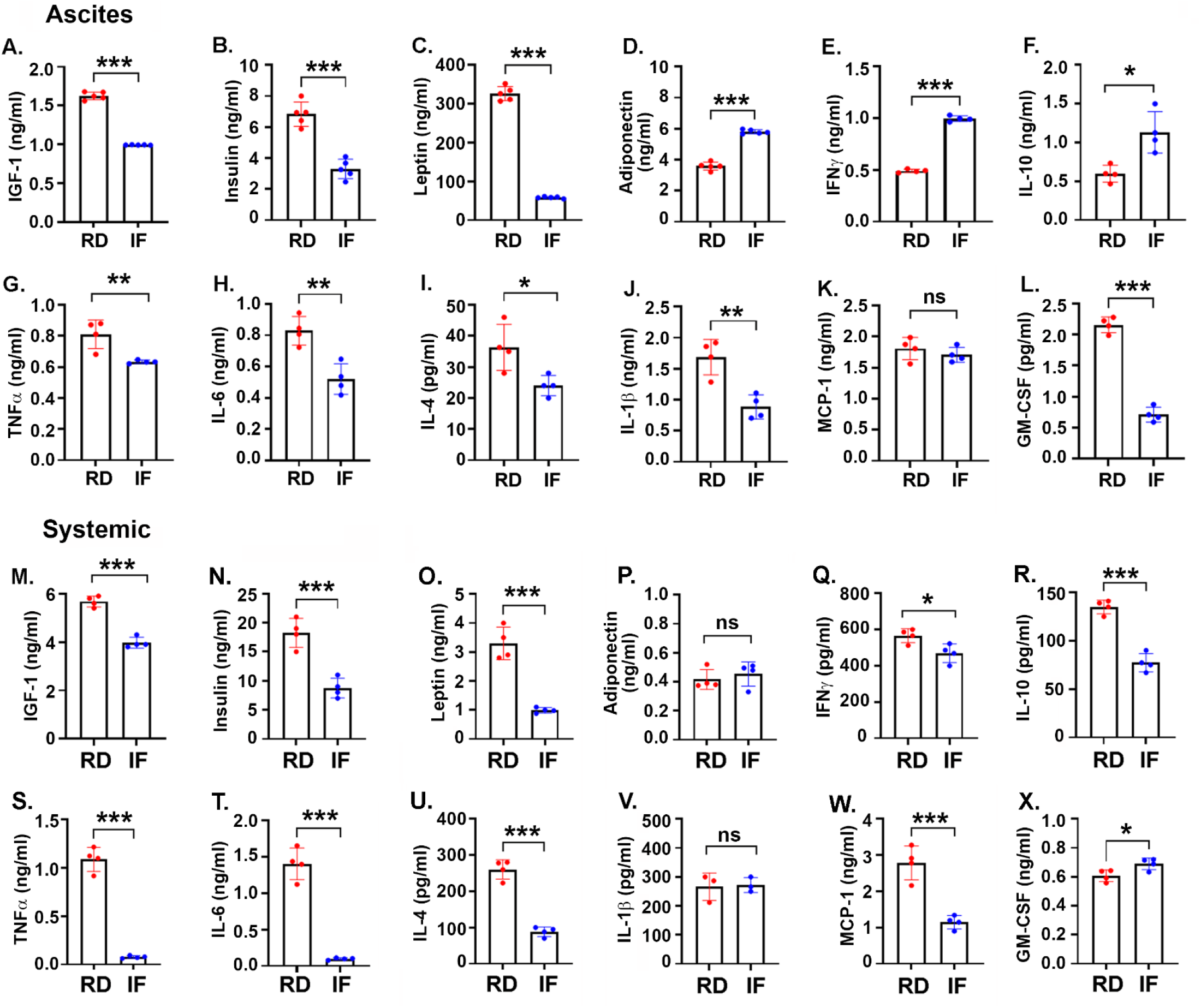
IF decreases tumor promoting growth and proinflammatory factors in EOC mouse models. Growth factors and pro inflammatory cytokines were measured by ELISA (n=4 or n=3) in the ascites (A-L) and plasma (M-X), collected at week 5 from ID8^p53-/-^ EOC bearing mice being subjected to RD or IF (n=4/group). (A, M) IGF-1, (B, N) Insulin, (C, O) Leptin, (D, P) Adiponectin, (E, Q) IFN γ, (F, R) IL-10, (G, S) TNFα, (H, T) IL-6, (I, U) IL-4, (J, V) IL-1β, (K, W) MCP-1 and (L, X) GMCSF. *p < 0.05, **p < 0.01, ***p < 0.001, ****P<0.0001, IF compared with RD group by Students T-test.

Thus, IF can differently modify and establish an anti-inflammatory and anti-growth milieu at both the tumor and systemic levels.

### IF remodels the antitumor T-cell immune response to EOC mouse tumors

Recent research has shown that short-term fasting or fasting mimicking diets can modify the immune response in a variety of disease models^38, 39^, but there are no reports in EOC. Immune profiling followed by tSNE projection was done in the tumor environment reflected by ascites and systemically reflected by blood obtained from ID8^p53-/-^ and ID8^p53-/-, PTEN-/-^ tumor carrying mice on IF and RD. The ascites of IF mice showed an increase in number of CD4^+^ and CD8^+^ T cells compared to RD mice (Fig. 3A-D, H; Fig S4). Analysis of the effector markers revealed that CD4^+^ T cells from IF mice displayed a robust Th1 phenotype, as evidenced by elevated intracellular IFNγ and decreased IL-4 expression (Fig. 3B, C, E-G; Fig S4). In contrast to RD mice, the effector, and cytotoxic markers of CD8^+^ T cell were significantly enhanced by IF as seen by increased intracellular production of IFNγ, granzyme B and perforin, (Fig. 3B, C, I-K; Fig S4). A similar pattern was observed in the blood of both models, with significant anti-tumor CD4 and CD8 T cell responses (Fig. S5). Thus, in preclinical models of EOC, our studies indicate that IF can stimulate an anti-tumor T cell response, which may lead to decreased tumor progression and enhanced survival.

**Fig 3:**
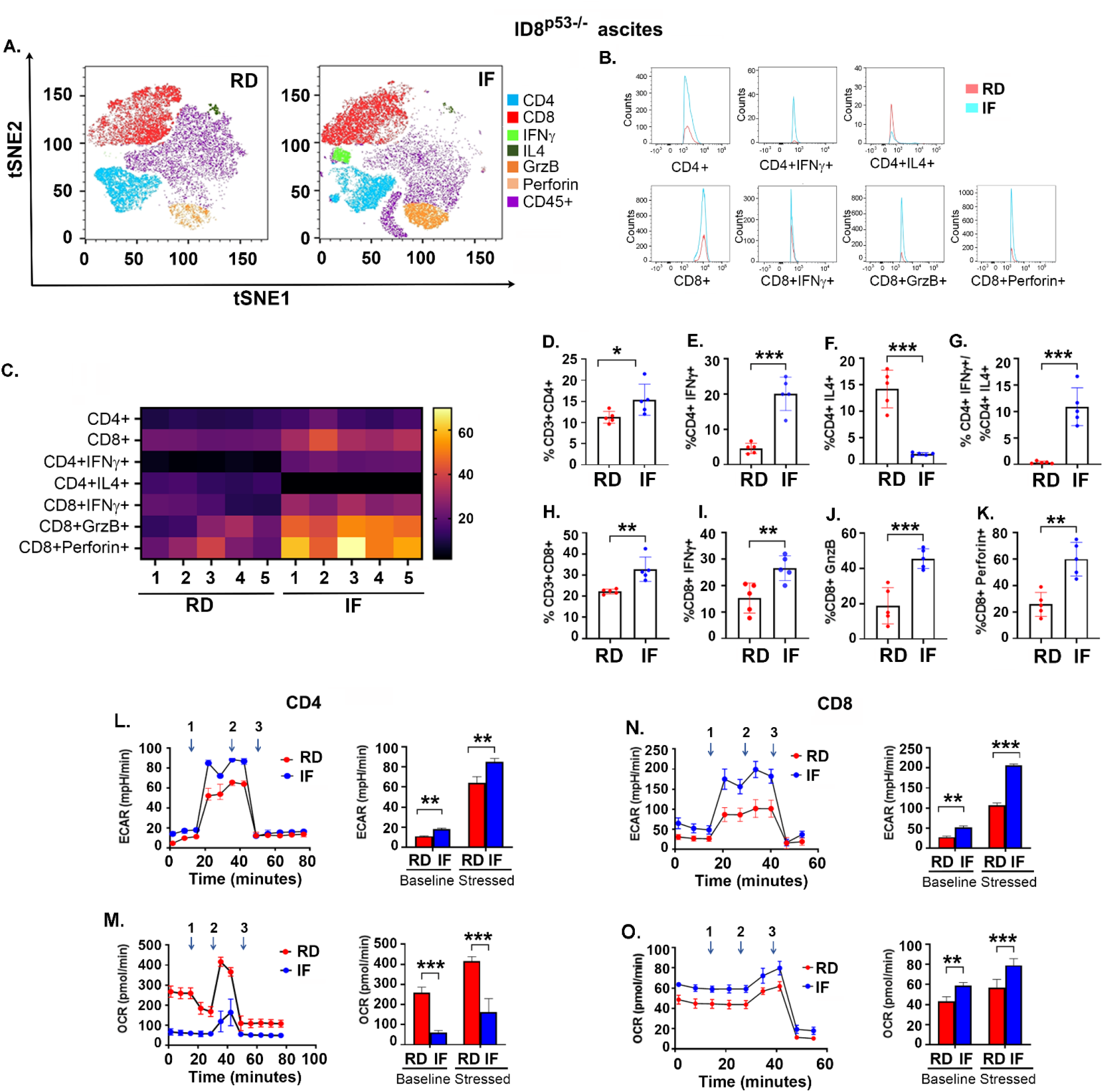
IF remodels the anti-tumor T-cell immune response to EOC mouse tumors. Female C57/BL6 mice were injected with ID8^p53-/-^ cells intraperitoneally and subjected to RD and IF, ascites and were collected at 5 weeks were analyzed by flow cytometry (n=5). (A) A t-SNE visualization of markers after gating on single, live, CD45+ CD3+, CD4+, CD8+, IFNγ, IL4, Granzyme B (Grz B) and perforin expression. (B) Histograms represent average counts of the main T cell subsets in RD and IF groups. (C) Heat map represents marker expression of the main T cell subsets in individual samples. Bar graph represents the percentage of T cell subsets (D) CD4+, (E) CD4+IFNγ+, (F) CD+IL4+, (G) ratio of CD4+ IFNγ+ to CD4+IL4+, (H) CD8+, (I) CD8+IFNγ+, (J) CD8+GrzB+ and (K) CD8+perforin+. CD4+ and CD8+ cells were isolated from spleens after 5 weeks of RD and IF mice and subjected to realtime XFe Seahorse analysis for bioenergetics profiling. Extracellular acidification rate (ECAR) an indicator of glycolysis was measured with port injections of (1) glucose, (2) oligomycin, and (3) 2-DG in (L) CD4+ cells and (N) CD8+ cells. Oxygen consumption rate (OCR) an indicator for mitochondrial respiration was assessed with port injections of (1) oligomycin, (2) FCCP, and a combination of (3) rotenone-antimycin given in (M) CD4+ and (O) CD8+ cells as described in methods. The bar graph represents basal and stressed OCR and ECAR (n=3). The experiment was repeated twice in two different sets of mouse experiment. *p < 0.05, **p < 0.01, ***p < 0.001, ****p < 0.0001, IF compared to RD group by Students T-test.

Function, differentiation, and activation of T cells are substantially regulated by metabolic modifications^40, 41^. However, modulation of T cell metabolism by IF is unknown. Compared to RD mice, splenic CD4^+^ cells from IF mice engaged in enhanced glycolysis, as evaluated by ECAR, and decreased OXPHOS, as measured by OCR^27^, determined by real-time bioenergetic profiling (Fig. 3L, M). Interestingly, splenic CD8^+^ T cells from IF mice were more active in both glycolysis and OXPHOS than CD8^+^ cells from RD mice (Fig. 3N, O). Consequently, CD4^+^ and CD8^+^ cells are metabolically more active in response to IF, which may facilitate their anti-tumor immune function.

Overall, IF can modulate the metabolism of CD4^+^ and CD8^+^ T cells, positioning them to execute their anti-tumor immune response.

### The absence of T cells reverses the anticancer effect of IF in EOC mouse tumors

To assess the importance of T cells in IF’s anti-tumor impact against EOC, we examined IF in nude mice devoid of thymus and with non-functional T cells^42^. Nude mice with ID8^p53-/-^ tumors that have been subjected to IF exhibited no change in survival compared to mice on RD (RD median survival of 39 days versus IF median survival of 38 days), body weight, abdominal circumference, or ascites volume (Fig. 4A-D). Measuring metabolic variables revealed that IF decreased IGF-1 while increasing insulin and having no effect on leptin and adiponectin (Fig. S4A). Some pro-inflammatory variables, such as IL-4, MCP-1, and GM-CSF, were reduced, while TNFα was elevated, and anti-inflammatory cytokines IFNγ and IL-10 were unaltered (Fig. S4B). Thus, while IF can partially alter the TME of EOC in nude mice, it cannot inhibit tumor growth, suggesting a key role for functioning T cells. To further confirm the dependence of IF on T cells in mediating its anticancer effect, we depleted CD4^+^and CD8^+^ cells in mice undergoing IF (Fig. S4C-F) and observed that CD8^+^ depletion reversed IF’s protective effect, while CD4^+^ depletion was less effective (Fig. 4E, S4G). In contrast to CD4^+^ depletion, CD8^+^ depletion reversed the IF mediated increase in survival and decrease in ascites and abdominal circumference (Fig. 4F, G and S4H, I). These findings imply that T cells are crucial mediators of IF’s anti-tumor action, with CD8^+^ cells playing a larger role than CD4^+^ cells.

**Fig 4:**
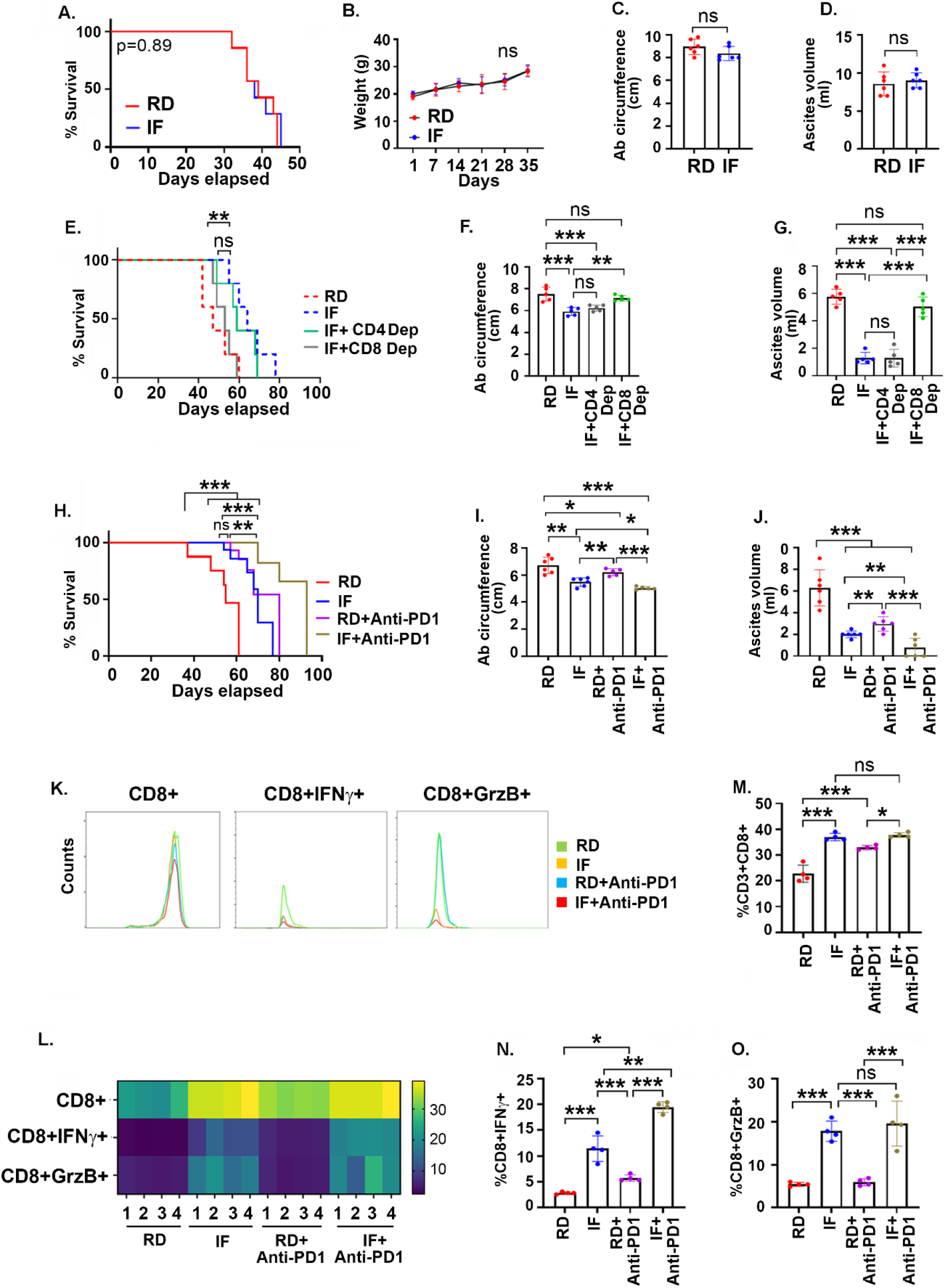
The absence of T cells reverses the anti-cancer effect of IF in EOC mouse tumors. ID8^p53-/-^ cells were injected intraperitoneally into Nude mice (Nu). After 1 week, mice were subjected to RD or IF (n=12/group) (A) Kaplan Meier graph indicating overall survival (n=12), p=0.89 by Gehan-Breslow-Wilcoxon test. (B) Average weekly body weight progression. Bar graph represents(C) average abdominal circumference, and (D) average ascites accumulated in RD and IF groups. Female C57/BL6 mice were injected with ID8^p53-/-^cells intraperitoneally and subjected to RD, IF, IF with CD4+ depletion, IF with CD8+ depletion and IF with IgG2b control. (E) Kaplan Meier graph indicating overall survival mice (n=12), p=0.38 IF compared to CD4 depletion, p= 0.009 IF compared to CD8 by Gehan-Breslow-Wilcoxon test. (F) Bar graph represents average ascites accumulated, and (G) Bar graph represents average abdominal circumference. Data with all control groups is shown in Fig. S6. Female C57/BL6 mice were injected with ID8^p53-/-^ cells intraperitoneally and subjected to RD, IF, anti-PD1 and IF combined with anti-PD1 treatments. Control RD and IF mice received IgG2b treatments. (H) Kaplan Meier graph indicating overall survival (n=12). p=0.006, IF compared to RD, p=0.0026 IF compared to IF and anti-PD1 combination, p=0.0006, RD compared to RD and anti-PD1 combination and p<0.0001 RD compared to IF and anti-PD1 combination by Gehan-Breslow-Wilcoxon test. Bar graph represents(I) average abdominal circumference, and (J) average ascites accumulated. Flow cytometry analysis was performed in ascites of all groups at 5 weeks. (K) Histograms representing average CD8+, CD8+IFNγ+ and CD8+GrzB+ abundance. (L) Heatmap representing individual frequency CD8+, CD8+IFNγ+ and CD8+GrzB+ cells. Bar graph representation of the percentages of (M) CD8+, (N) CD8+IFNγ+ and (O) CD8+GrzB+ in RD, IF, anti-PD1 and IF combined with anti-PD1 treated mice. *p < 0.05, **p < 0.01, ***p < 0.001, ****p<0.0001, by one way ANOVA, followed by Sidak multiple comparison test.

Use of monoclonal antibodies to block the interaction between programmed cell death protein (PD-1) and its counter ligand programmed death ligand 1 has demonstrated promising therapeutic effectiveness in various malignancies, except in EOC^43, 44^. Since IF improved T cell response, we examined whether it could also improve immunotherapy outcomes. ID8^p53-/-^ injected mice treated with anti-PD 1 in combination with IF exhibited a substantial increase in survival, with a median survival of 93 days compared to 55 days for the RD mice, 70 days for mice on IF alone and 80 days for mice treated with anti-PD1 alone (Fig. 4H). Lower abdominal circumference and ascites volume represented the decreased tumor load in the combination group compared to the single therapy and IF groups (Fig. 4I, J). In the ascites, the anti-tumor CD8^+^ T cell response was enhanced in the combination group, as seen by an increase in the CD8^+^ T cells expressing IFNγ and granzyme B, although IF alone was equally efficient in increasing CD8^+^ cell number and granzyme B expression (Fig. 4K-O). An identical improvement in CD8^+^ T cell response was also detected at the systemic level (Fig. S6J-N).

These findings demonstrate that T cells are a critical component of IF’s anticancer action, and that IF can enhance the anti-tumor CD8^+^ T response induced by PD-1 inhibition.

### IF induces ketogenesis

As any dietary intervention would result in central metabolism reconfiguration, we performed untargeted metabolomics on plasma from EOC mice on RD and IF to determine the molecular metabolic mediators of IF (Fig. S7A). Principal component analysis^29, 30^ demonstrated strong separation along plasma metabolites of RD and IF mice (Fig. S7B). The heatmap representation of the significantly 471 altered metabolites revealed clear alterations in all the key super-metabolism pathways (Fig. 5A). In response to IF, the Metaboanalyst Pathway enrichment analysis^30^ identified ketone body metabolism as the most significantly enriched pathway (p≤0.041, FDR≤0.05) (Fig. 5B, S7C). Metabolite concentrations of the ketone bodies acetoacetate and BHB were significantly increased (Fig. 5C, D). In another group of mice, we further validated the increased levels of BHB, the major ketone body generated in the liver and transported to other tissues to be used as fuel^45^, in plasma and ascites of fasting mice as detected by LC/MS/MS (Fig. 5E, F) and ELISA (Fig. 5G, H). Enhanced expression of liver ketogenic enzymes, including ACAT, HMGCS2, HMGCL, and BDH1, correlated with elevated BHB levels (Fig. 5I, J), indicating enhanced ketogenesis in the liver.

**Fig 5:**
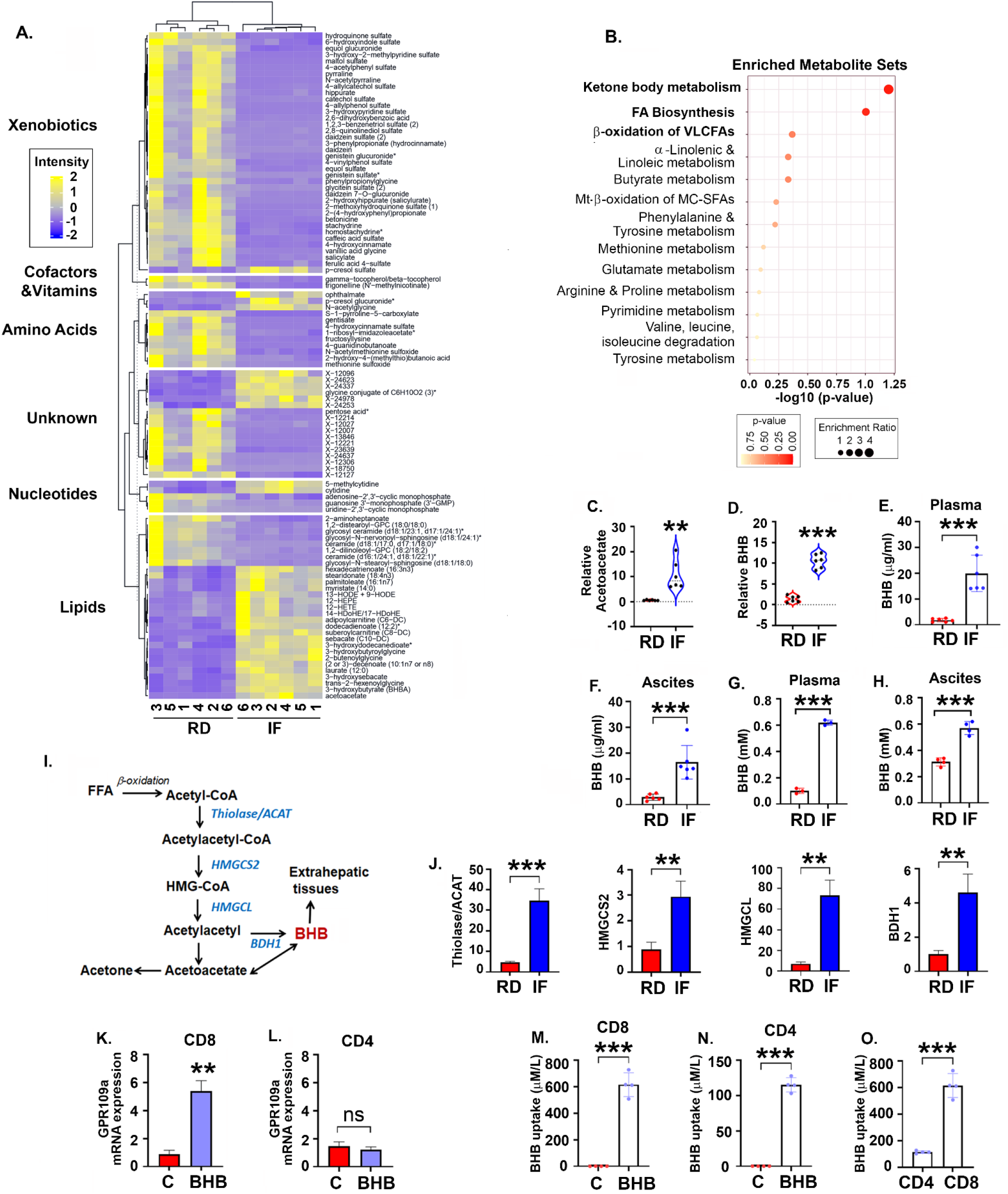
IF induces Ketogenesis: Untargeted metabolomics was performed in the plasma from ID8^p53-/-^ tumor bearing mice subjected to RD or IF at 5 weeks (n=6/group). (A) Heatmap representation of the significantly 471 altered metabolites in the major super-metabolism pathways. (B) Pathway enrichment analysis (p < 0.05; FDR < 0.05). Scaled metabolite intensity graphs showing levels of (C) acetoacetate and (D) BHB detected during untargeted metabolomics. BHB levels were measured in plasma (E) and (F) ascites using LC-MS/MS (n=6/group). BHB levels were measured in plasma (G) and (H) ascites by ELISA (n=5/group). (I) Pathway showing enzymes involved in ketogenesis. (J) mRNA expression of liver ketogenic enzymes (J) ACAT, HMGCS2, HMGCL and BDH1 measured by RT-PCR. (K) mRNA expression of GPR109a measured by RT-PCR on naïve (K) CD8+ and (L) CD4+ cells in absence or presence of 10mM BHB. Uptake of BHB levels were measured using LC-MS/MS in naïve (M) CD8+ cells and (N) CD4+ cells after 72h exposure to BHB. (O) Ratio of intracellular BHB levels taken up by CD4+ and CD8+ cells. **p < 0.01, ***p < 0.001, ****p<0.0001, IF compared to RD and BHB compared to RD assessed by Student t tests.

To determine if the bioactive BHB can communicate with and impact T cells, the expression of the BHB receptor, GPR109a^46, 47^, was evaluated on CD4^+^ and CD8^+^ T cells isolated from naïve mice and treated with BHB. GPR109a was expressed at a baseline level in both CD4^+^ and CD8^+^ cells, whereas BHB treatment increased GPR109a expression in CD8^+^ T cells, but not in CD4^+^ T cells (Fig. 5K, L). To further validate, we assessed the BHB uptake by CD4^+^ and CD8^+^ T cells following BHB exposure. Both CD4^+^ and CD8^+^ cells were able to uptake BHB (Fig. 5M, N), but CD8^+^ cells were able to uptake almost 5 times more BHB than CD4^+^ T cells (Fig. 5O).

Overall, IF generated substantial metabolic remodeling at the systemic and cellular levels, with a rise in ketone bodies and metabolism that may have a more profound effect on CD8^+^ T cell activity.

### BHB inhibits tumor growth and promotes antitumor CD8^+^ T cell responses in EOC mouse models

To determine if BHB can replicate the anti-tumor and increase T cell response of IF, we administered BHB to mice with ID8^p53-/-^ and ID8^p53-/-PTEN-/-^ tumors. BHB treatment significantly enhanced overall survival in mice with ID8^p53-/-^ tumors (median survival 63 days versus 50 days) and ID8^p53-/-PTEN-/-^ tumors (median survival 70.5 days versus 53.5 days) (Fig. 6A, S8A). The body weights of BHB treated mice increased more slowly than those of RD mice, whereas abdominal circumference and ascites volume were significantly reduced in both models (Fig. 6B-D, S8B,C) compared to control. Profiling of the T cell response revealed an unexpected decrease in CD4^+^ T cells, although intracellular IFNγ levels still rose (Fig. 6E-I, S8E, G, H). BHB treated ID8^p53-/-^ mice exhibited an increase in IL-4 that resulted in a non-significant Th1 response of CD4^+^ cells (Fig. 6J, K), but BHB suppressed IL4 production in the ID8^p53-/-PTEN-/-^ mice (Fig S8I). BHB induced a robust increase in CD8^+^ T cell number and effector markers IFNγ and Granzyme B (Fig. 6E-G, L-N, S8J-L), as well an increase in CD8:CD4 ratio (Fig. 6O). In the blood of both models, a comparable systemic profile of increased T cell response was also seen in response to BHB (Fig. S9, S7F, M-R, data shown for ID8^p53-/-^).

**Fig 6:**
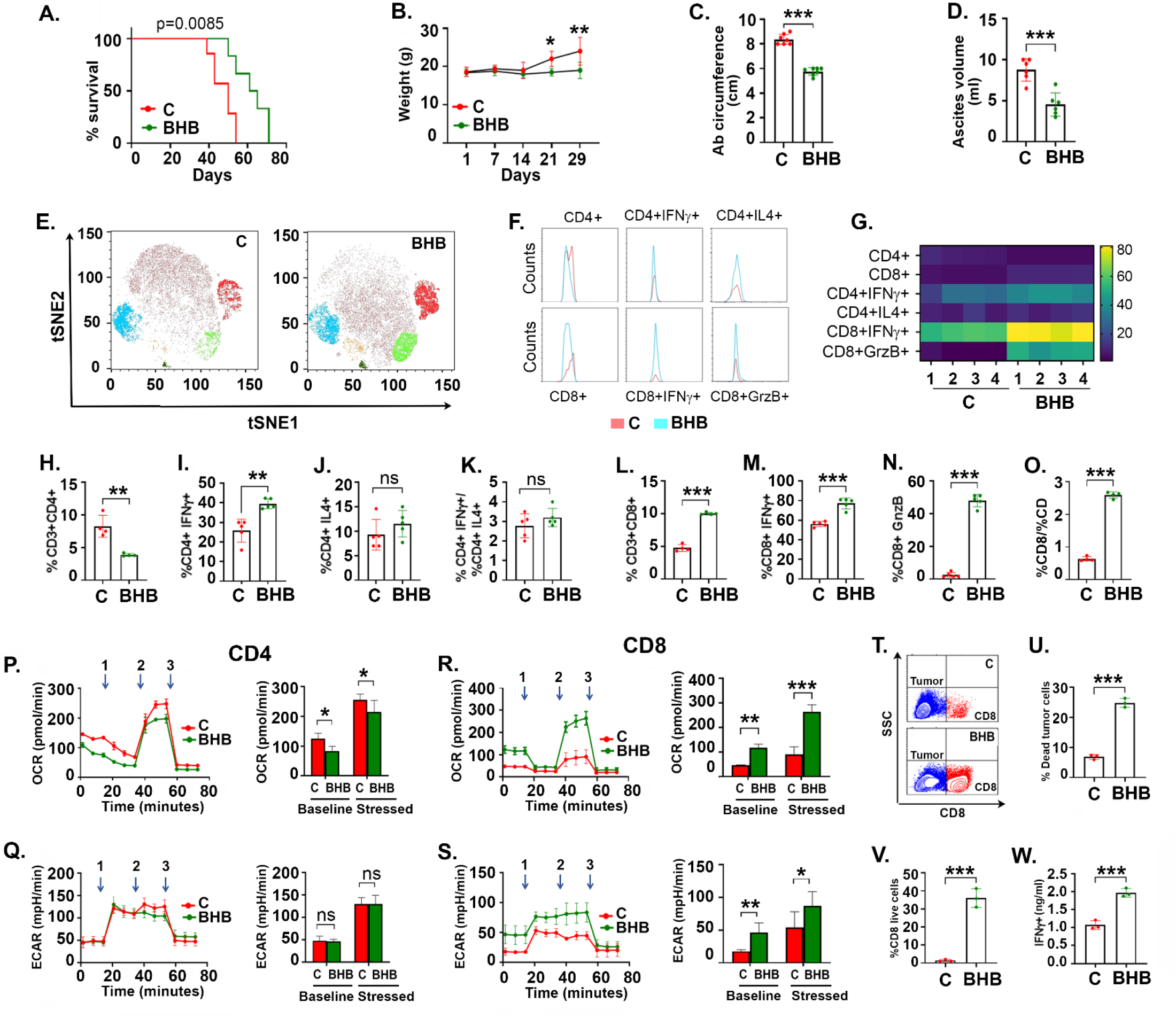
BHB inhibits tumor growth and promotes anti-tumor CD8+ T cell responses in EOC mouse models. ID8^p53-/-^ cells were injected intraperitoneally into femaleC57/B6 mice. After 1 week, mice were treated with either BHB (300mg/kg bd wt x 3 doses/week) or vehicle PBS (control mice, C). (A) Kaplan Meier graph indicating overall survival (n=12), p=0.0085 by Gehan-Breslow-Wilcox test. (B) Average weekly body weight progression. Bar graph represents(C) average abdominal circumference, and (D) average ascites volume. Immune profiling was performed at 5 weeks (n=4). (E) A t-SNE visualization of markers after gating on single, live, CD45+ CD3+, CD4+, CD8+, IFNγ+, IL4+ and Granzyme B + (GrzB) cells. (F) Histogram represents counts of the T cell subsets in ascites of C and BHB treated mice. (G) Heat map represents marker expression of the T cell subsets. Bar graph represents the percentage of T cell subsets (H) CD4+, (I) CD4+IFNγ+, (J) CD+IL4+, (K) ratio of CD4+ IFNγ+ to CD4+IL4+, (L) CD8+, (M) CD8+IFNγ+, (N) CD8+Grz B+ and (O) ratio of CD8+ to CD4+ cells. CD4+ and CD8+ cells isolated from spleens were subjected to real-time XFe Seahorse analysis for bioenergetics profiling. Oxygen consumption rate (OCR) an indicator for mitochondrial respiration was assessed with port injections of (1) oligomycin, (2) FCCP, and a combination of (3) rotenone-antimycin, in (P) CD4+ and (R) CD8+ cells as described in methods. Extracellular acidification rate (ECAR) an indicator of glycolysis was measured with port injections of (1) glucose, (2) oligomycin, and (3) 2-DG in (Q) CD4+ cells and (S) CD8+ cells. The bar graph represents basal and stressed OCR and ECAR (n=3). The experiment was repeated twice in two different sets of mouse experiment (n=3). Splenic naïve CD8+ cells after activation were treated or untreated with 10mM BHB and co-cultured with 7AAD labeled ID8^p53-/-^ cells and analyzed for percentage of live and dead ID8^p53-/-^ and CD8+ cells (T). (U) Bar graph represents percentage of dead ID8^p53-/-^ tumor cells. (V) Bar graph represents percentage of CD8+ cells. (W) IFNγ levels measured in supernatant collected from co-culture by ELISA. *p < 0.05, **p < 0.01, ***p < 0.001, ****p<0.0001, BHB compared to C, assessed by Student t tests.

Bioenergetic profiling of CD4^+^ and CD8^+^ cells from the spleens of control and BHB treated mice revealed a significant increase in glycolysis measured as ECAR and OXPHOS measured as OCR^23, 27, 30^ of CD8^+^ T cells alone (Fig 6R,S), but CD4^+^ bioenergetics were relatively unaffected, with increase in OCR alone (Fig 6P,Q). Activated splenic CD8^+^ and CD4^+^ cells were co-cultured with 7AAD labeled ID8^p53-/-^ tumor cells in absence or presence of BHB to confirm that the increased effector expression and metabolic capabilities translated to an increase in function. BHB treated CD8^+^ cells caused more tumor cell death than untreated CD8^+^ cells (Fig 6T, U). Intriguingly, BHB treated CD8^+^ also demonstrated increased proliferation and IFNγ levels (Fig 6V, W), indicating enhanced antitumor capability. In contrast, BHB treated CD4^+^ were unable to induce tumor cell death, despite enhanced proliferation and a non-significant increase in IFNγ levels (Fig S9J-M).

Thus, BHB is an essential modulator of the anti-tumor CD8^+^ response observed during IF in EOC preclinical models.

### IF improves EOC survival and antitumor immune response better than BHB

We directly compared the outcomes of IF and BHB in the ID8^p53-/-^ model due to the observation that IF appears to impact both CD4^+^ and CD8^+^ cells, as well as the central metabolism. While BHB treatment enhanced survival compared to untreated control mice on RD, the IF group showed a greater gain in survival (RD median survival 41 days versus 56 days in BHB versus 65 days in IF) (Fig. 7A). Weight changes were comparable across BHB treatment and IF (Fig. 7B). IF mice exhibited lower abdominal circumference and lower ascites volume than BHB mice (Fig. 7C, D). Similar decreases in insulin, IGF-1, and leptin levels were observed in both groups, although adiponectin levels were unaffected by BHB in the ascites (Fig. 7E). Both raised the anti-inflammatory IFNγ, although the increase in IF mice was greater (Fig. 7F). All other pro-inflammatory cytokines including TNFα, IL-6, MCP-1 and GM-CSF were reduced in both BHB and IF, although IL-4 and IL-1β were unaffected by BHB in the ascites (Fig. 7G-L). Immune profiling of ascites demonstrated that IF significantly increased CD4^+^ and CD8^+^ cell populations, whereas BHB significantly increased only CD8^+^ cells and a trend towards decrease in CD4^+^ cells as seen before (Fig. 7M-P, S). IF demonstrated Th1 intracellular indicators of increased IFNγ and decreased IL-4 in CD4^+^, whereas BHB had no effect on IL-4 as before (Fig. 7M-O, Q, R). Both IF and BHB increased intracellular IFNγ and GrzB in the CD8^+^ cells, although IF caused a greater proportional rise (Fig. 7T, U). At the systemic level, a comparable T cell based immunological response was observed (Fig. S10), with differences in CD4^+^IFNγ^+^ and CD8^+^ Granzyme B^+^ stimulation by BHB.

**Fig 7:**
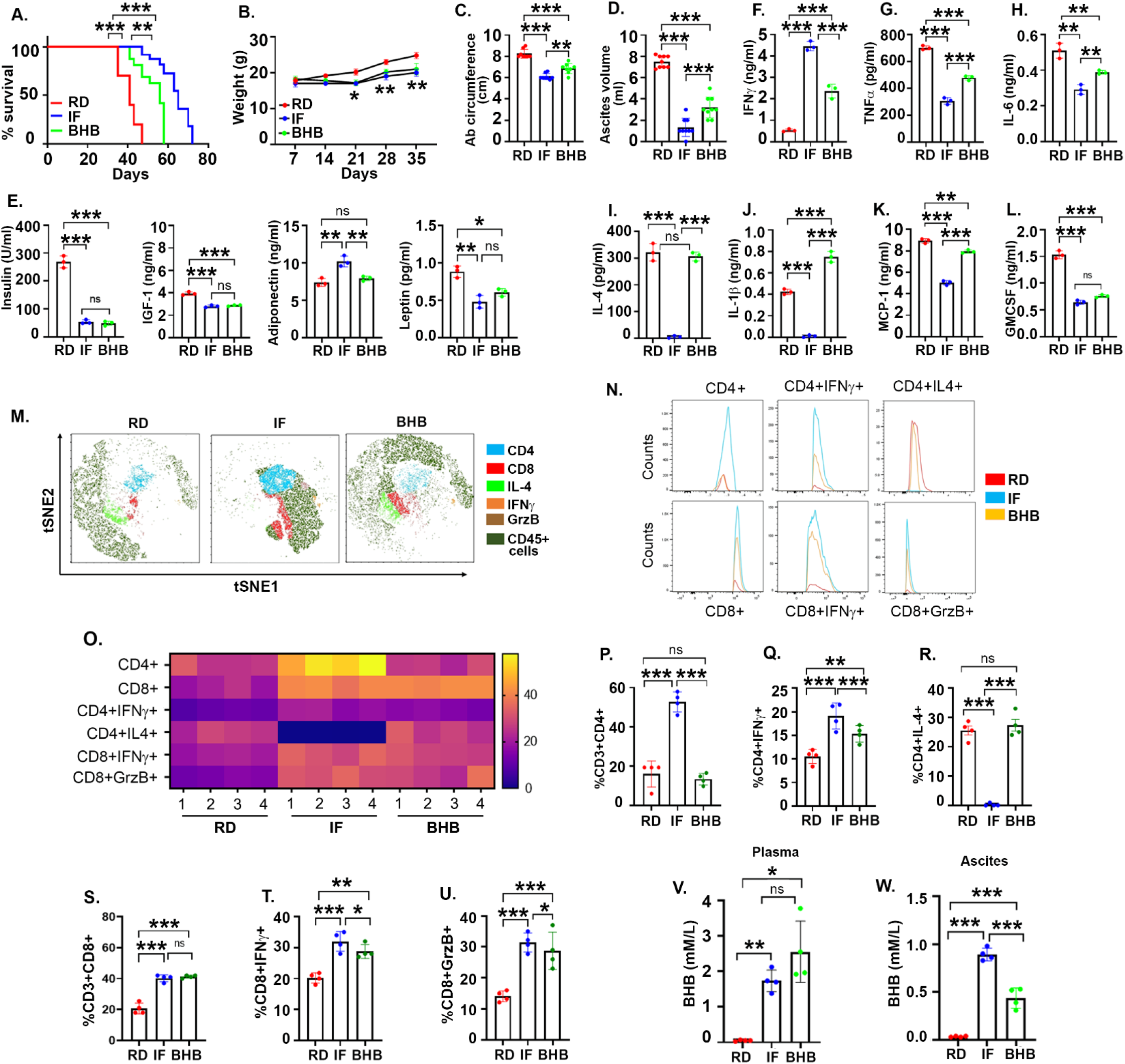
IF improves EOC survival and anti-tumor immune response better than BHB. ID8^p53-/-^ cells were injected intraperitoneally into C57/B6 female mice. After 1 week, mice were subjected to control (C, untreated), IF or BHB treatments. (A) Kaplan Meier graph depicting overall survival (n=12/group), p<0.0001 IF compared to RD, p<0.0001 BHB compared to RD and p=0.0016 BHB compared to IF by Gehan-Breslow-Wilcoxon test. (B) Average weekly body weight progression. Bar graph represents (C) average abdominal circumference and (D) average ascites volume. Growth factors and cytokines were measured in ascites collected at week 5 by ELISA (n=3/group), (E) Insulin, IGF-1, leptin, and adiponectin, (F) IFNγ, (G) TNFα, (H) IL-6, (I) IL-4, (J) IL-1β, (K) MCP-1 and (L) GMCSF. Immune profiling was performed after 5 weeks of treatment (n=4/group). (M) A t-SNE visualization of markers after gating on single, live, CD45+ CD3+, CD4+, CD8+, IFNγ+, IL-4+ and Granzyme B+ (GrzB) cells. (N) Histogram represents average counts of the T cell subsets measured. (O) Heat map represents marker expression of the measured T cell subsets in individual samples. Bar graph represents the percentages of T cell subsets (P) CD4+, (Q) CD4+IFNγ+, (R) CD4+IL4+, (S) CD8+, (T) CD8+IFNγ+, and (U) CD8+Gnz B+. BHB levels were measured in (V) plasma and (W) ascites of C, IF and BHB mice. *p < 0.05, **p< 0.01, ***p < 0.001, ****p<0.0001, by one way ANOVA, followed by Sidak multiple comparison test.

To determine whether the accumulation of BHB differed between the two approaches, we measured BHB levels in the plasma and ascites of mice from each group. BHB therapies resulted in greater BHB levels in circulation (Fig. 7V). At the tumor microenvironment level of the ascites, IF resulted in significantly higher BHB levels than BHB treatments (Fig. 7W).

These observations indicate that IF and BHB have nearly similar impact on the immune response and inflammatory factors, suggesting that other physiologic metabolic alterations or immune cells may be responsible for the greater anti-cancer activity of IF in preclinical models of EOC.

## Discussion

Intermittent fasting (IF) is a dietary-based interventional therapy that alternates between fasting and eating over time. The benefits of IF in preventing a variety of diseases and increasing lifespan have prompted interest in its benefits in the field of cancer^13^. In a number of tumors outside EOC and gynecologic cancers, numerous studies have examined various IF regimens with varying results. Here, we emphasize IF’s capacity to induce anti-tumor immunostimulatory effects via its metabolic mediator BHB, which can be leveraged to limit EOC and increase patient survival.

In preclinical models of EOC with increasing aggressiveness^21^, the IF regimen comprising a 16-hour fast and an 8-hour feeding window for 5 days per week significantly increased survival. The enhanced survival and decreased tumor growth were associated with an enhanced T cell response against the tumors characterized by a rise in the percentage of CD4^+^ and CD8^+^ cells, an enhancement of their effector function and modulation of their metabolic activity. While it has been shown that a restrictive diet exerts its anti-tumor effects via a decrease in metabolic and growth factors such as IGF-1 and insulin, direct differential stress on tumor cells, nutritional restriction, and reduced inflammation; recent studies have highlighted their immunomodulatory effect^7, 9^. Our work demonstrate that T cells were important for IF-mediated enhanced survival and decreased tumor growth, since nude mice with ovarian cancer gained no benefit from IF. We also found that IF enhanced the efficacy of anti-PD1 immunotherapy in preclinical models of EOC. Studies on T cell depletion reveal that CD8^+^ T cells are more crucial than CD4^+^ T cells in IF-induced anti-tumor T cell response. The correlation between higher effector T cell function and increased metabolic activity of T cells in response to IF suggests that reprogramming of cellular metabolism may be a potential mechanism by which IF regulates the number and function of T cells. Together, our research reveals that IF’s immunomodulatory capacity is the primary reason of its anti-tumor activity in preclinical models of EOC.

IF or fasting alone has been studied in a variety of malignancies, using different fasting duration and diet type^13^. The physiologic and metabolic alterations differ based on duration and frequency of fasting, which may determine the outcome and the underlying mechanisms. The antitumor effects of IF have been attributed to weight-loss, creating a tumor environment characterized by decreased growth and metabolic factors, inflammation, and oxidative stress. Such changes can induce autophagy, apoptosis and enhance chemotherapy response by differentially protecting normal cells but not the cancer cells. More recently, such dietary interventions have also been shown to promote anti-tumor immune responses^7, 48^. Studies combining various fasting schedules with chemotherapy have demonstrated a significant slowing of tumor progression in preclinical models of many cancer types^15, 49^. While long-term fasting schedules have not been evaluated in cancer patients, a handful of small clinical trials have evaluated the combination of short-term fasting (24-72h fast) with chemotherapy, with an emphasis on lowering chemotherapy side effects and enhancing QoL measures^16, 17,18^. Collectively, these studies demonstrate that fasting in combination with chemotherapy in cancer patients, including ovarian cancer patients, is achievable with a moderate to high adherence rate and results in a substantial enhancement of QoL. Current and future clinical trials are being actively performed and designed to also investigate the mechanisms and effect on cancer cells and patient survival. Thus, IF is a potential promising approach that can be incorporated into patient treatment and management to not only improve QoL, but also successfully inhibit tumor growth, enhance chemotherapy and boost anti-cancer immune response.

Ketone bodies are generated predominantly in the liver by ketogenesis from beta-oxidation of free fatty acids (FFA) and then transported to extrahepatic peripheral organs for oxidation, a process known as ketosis or ketolysis. Under many physiologic situations, including caloric restriction, fasting, starvation, post-exercise, pregnancy, neonatal stage, and low-carbohydrate diets such as the ketogenic diet, ketone body oxidation becomes the primary alternative contributor to energy metabolism in extrahepatic organs, such as muscle, heart, and brain. To determine the metabolic determinant of IF’s immunomodulatory function we performed plasma metabolomic profiling. IF triggered dramatic alterations in the central metabolism, including the metabolic switch from glucose to fatty acids and ketone bodies as the primary energy source. This was demonstrated by the enhancement of ketone body metabolism in the plasma of IF-fed mice, the considerable increase in systemic BHB levels, and the enhancement of ketogenesis enzyme expression in the liver. Elevated amounts of BHB were similarly detected in the TME of ascites. BHB is a potent endogenous metabolite that functions as a signaling mediator, epigenetic regulator, protein post-translation modifier, and regulator of inflammation and oxidative stress, in addition to its role as an alternative energy source, ^45, 50, 51^.

With the recent trials of ketogenic diets in cancer, the role of ketosis in cancer biology is better understood; nevertheless, BHB-based research has yielded contradictory results. Studies have suggested that BHB supplementation could inhibit, stimulate, or have no effect on tumor growth^12, 52, 53, 54, 55, 56,47, 57^. However, *the effect of BHB in ovarian cancer is largely unknown.* A recent *in vitro* study demonstrated that BHB had no influence on the proliferation or EMT or stem cell characteristics of ovarian cancer cell lines in a low glucose environment, despite an increase in few enzymes of the ketogenesis pathway^58^. Few recent studies indicate modest benefits of ketogenic diet for ovarian cancer patients^59, 60^. Recent research demonstrates that BHB has an immunomodulatory effect that can enhance the immune response against tumors. By boosting CD8^+^ T cells, BHB has been shown to improve the efficacy of checkpoint-based immunotherapy in melanoma, lung, and renal cell cancer models^11^. In healthy human volunteers, BHB has been demonstrated to have a dramatic effect on CD4^+^ and CD8^+^ T response and improve T cell immunity^61^. Our study indicates that treatment with BHB replicated tumor regression, enhanced survival, modulation of metabolic growth factors and cytokine, and immunomodulatory effects of IF. Interestingly, BHB had a greater immunostimulatory effect on CD8^+^ cells than on CD4^+^ cells, as indicated by substantial increase in CD8^+^ T cell numbers and effector markers IFNγ and granzyme B whereas its effect on CD4^+^ cells was minimal. This differential effect may be explained by the higher expression of the BHB receptor GPR109a^46^, and the increased uptake of BHB by CD8^+^ cells compared to CD4^+^T cells. Compared to CD4^+^ cells, the BHB-treated mice’s CD8^+^ cells exhibited significantly higher bioenergetics, as seen by an increase in OCR and ECAR. Functionally, BHB-treated CD8^+^ cells were enhanced in their ability to kill tumor cells, as seen by an increase in CD8^+^ T cells and IFNγ relative to CD4^+^ cells, demonstrating that the anti-tumor effect of BHB is a direct result of its effect on the function and metabolism of CD8^+^ T cells. Overall, BHB appears to be a promising therapeutic agent, which may open new avenues for its application in the treatment and prevention of EOC.

A direct comparison of IF and BHB found that IF is more effective than BHB in slowing tumor progression and prolonging survival. The only significant difference between IF and BHB profiles was the decreased IL-4 and enhanced CD4^+^ activation. However, the observation that depletion of CD8^+^ reversed the anti-tumor impact of IF, whilst CD4^+^ depletion had only a partial effect, suggests a stronger role for the CD8^+^ T-cell based immune response in IF too, but one that may require equal activation of CD4^+^ cells to reach its full potential. Recent studies demonstrate the significance of CD4^+^ T cells in the cytotoxic effector activity of CD8^+^ T^62^ cells. It is possible that the greater effect of IF is mediated by other metabolic alterations or another anti-tumor metabolite(s) that inhibits tumor-promoting machinery, hence establishing an anti-tumor milieu in the host. It is also known that fasting induces widespread changes in the gut microbiome that are disease-alleviating and anti-cancer^63^.

Individual metabolic heterogeneity and adherence variability make it difficult to regulate ideal circulation BHB levels with food, which makes the therapeutic application of BHB appealing. However, the contradictory reports show that BHB plays a paradoxical role. Many parameters may be contributing to the diverging effects, such as the nutritional and bioenergetic state of the tumor cells, the right threshold levels of BHB, or the tumor cells’ ability to undergo ketolysis, as well as the tumor’s type and its location. Those parameters may determine whether BHB is used as a fuel by tumor cells to live and proliferate, is preferentially taken up by immune cells for immunological activation, or influences transcription gene expression as an epigenetic regulator. Therefore, more research is required to investigate BHB as a potential therapeutic agent and to determine the optimal balance between its effects on tumor cells and immune cell activity. On the other hand, findings regarding the anti-tumor effect of IF have been promising, indicating powerful anti-cancer benefits, and no evidence of increasing cancer progression. While we have detailed the immune-enhancing benefits of dietary alterations, there is evidence that dietary modifications also involve metabolic, microbiome, and genetic epigenetic changes, and it is the interaction of those factors that results in tumor suppression.

In conclusion, strategies based on fasting show great promise and our data supports the study of IF in pilot studies of EOC patients. However, there is a clear need to standardize the restriction length and timings as well as to monitor for any negative effects, particularly on immune cells and tumor cells.

## Notes

**Funding:** This work was supported by R01CA249188 awarded to RR and HFCI postdoctoral fellowship to MPU and HS. The funding source had no part in determining the study design or submission of the manuscript.

**Conflict of Interest:** We have no conflict or potential competing conflicts. All authors have seen and approved the manuscript, and that it hasn’t been accepted or published elsewhere.

### Competing Interest Statement

The authors have declared no competing interest.

