## Supplemental Material for "Intermittent Fasting induced ketogenesis inhibits mouse epithelial ovarian tumors by promoting anti-tumor T cell response"

### SUPPLEMENTARY TABLES

#### Table S1: List of antibodies used in vivo

| Antibody | Source | Identifier |
|---|---|---|
| *InVivo*Mab anti-mouse PD-1 | BioXCell | BE0146 |
| *InVivo*Mab rat IgG2b isotype | BioXCell | BE0090 |
| *InVivo*Plus anti-mouse CD4 | BioXCell | BP0003 |
| *InVivo*Mab anti-mouse CD8α | BioXCell | BE0061 |

#### Table S2: List of fluorochromes for flowcytometry analysis

| Antibody | Clone | Fluorochrome | Source |
|---|---|---|---|
| CD3 | 17A2 | AF700 | Biolegend |
| GRANZYME B | QA16AO2 | PE-CY5 | Biolegend |
| PERFORIN | S16009A | PE | Biolegend |
| CD4 | GK.1.5 | BV510 | Biolegend |
| IFN-GAMMA | XMG1.2 | PE-/DAZZLE 594 | Biolegend |
| CD45 | 30-F11 | PE-CY5.5 | Biolegend |
| IL-4 | 11B11 | PE-CY7 | Biolegend |
| CD8 | 53-6.7 | PERCP-CY5.5 | Biolegend |

#### Table S3: List of ELISA kits

| ELISA | Source | Identifier |
|---|---|---|
| IL-6 | Biolegend | 431301 |
| IFN γ | Biolegend | 430801 |
| IL-1 β | Biolegend | 432604 |
| MCP 1 | Biolegend | 432704 |
| TNF α | Biolegend | 430901 |
| IL 4 | Biolegend | 431104 |
| GMCSF | Biolegend | 432204 |
| IL 10 | Biolegend | 431414 |
| Insulin | Cayman Chemical | 589501 |
| IGF-1 | R and D Biosystems | MG100 |
| Leptin | R and D Biosystems | MOB00B |
| Adiponectin | R and D Biosystems | MRP300 |
| Albumin | Cayman Chemical | 500840 |
| Beta hydroxy butyric acid | Cayman Chemical | 700190 |

#### Table S4: Adipokine array

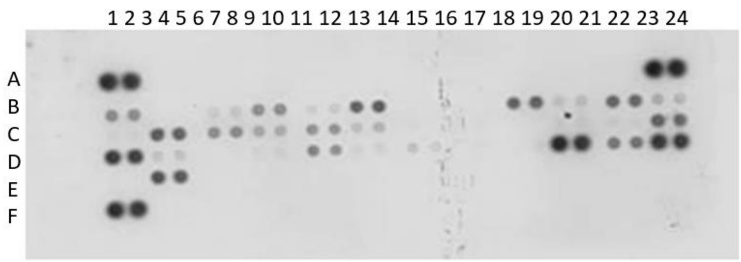


| Coordinate | Analyte |
|---|---|
| A1, A2 | Reference spots |
| A23, A24 | Reference spots |
| B1, B2 | Adiponectin |
| B3, B4 | AgRP |
| B5, B6 | ANGPT-L3 |
| B7, B8 | C-Reactive protein |
| B9, B10 | DPPIV |
| B11, B12 | Endocan |
| B13, B14 | Fetuin A |
| B15, B16 | FGF acidic |
| B17, B18 | FGF 21 |
| B19, B20 | HGF |
| B21, B22 | ICAM-1 |
| B23, B24 | IGF-1 |
| C1, C2 | IGF-11 |
| C3, C4 | IGFBP-1 |
| C5, C6 | IGFBP-2 |
| C7, C8 | IGFBP-3 |
| C9, C10 | IGFBP-5 |
| C11, C12 | IGFBP-6 |
| C13, C14 | IL-6 |
| C15, C16 | IL-10 |
| C17, C18 | IL-11 |
| C19, C20 | Leptin |
| C21, C22 | LIF |
| C23, C24 | Lipocalin-2 |
| D1, D2 | MCP-1 |
| D3, D4 | M-CSF |
| D5, D6 | Oncostatin M |
| D7, D8 | Pentraxin 2 |
| D9, D10 | Pentraxin 3 |
| D11, D12 | Pref-1 |
| D13, D14 | RAGE |
| D15, D16 | RANTES |
| D17, D18 | RBP4 |
| D19, D20 | Resistin |
| D21, D22 | Serpin E1 |
| D23, D24 | TIMP-1 |
| E1, E2 | TNF-a |

| | |
|---|---|
| E3, E4 | VEGF |
| F1, F2 | Reference spots |
| F23, F24 | PBS (Negative control) |

**Table S5: List of Primers**

| | |
|---|---|
| BDH 1 -F | 5'-GAA AGT GGT GGA GAT TGT CCG C-3' |
| BDH1 R | 5'-TGT AGG TCT CCA GGC TGG TGA A-3' |
| GPR 109a F | 5′-CGAGGTGGCTGAGGCTGGAATTGGGT-3 |
| GPR 109a R | 5′-ATTTGCAGGGCCATTCTGGAT-3′ |
| ACAT F | 5'-TGA GAG CAC CTC CAG AAC AAG G-3' |
| ACAT R | 5'-GGA CGA ATA GGA TGA GGA GTG C-3' |
| HMGCS2 F | 5'-CCT TGA ACG AGT GGA TGA GA-3' |
| HMGCS2 R | 5'-CAG ATG CTG TTT GGG TAG CA-3' |
| HMGCL F | 5’-ACCACCAGCTTTGTGTCTCC-3’ |
| HMGCL R | 5′-GAGGCAGCTCCAAAGATGAC-3′ |
| L27-F | 5'-ACA TTG ACG ATG GCA CCT C-3' |
| L27-F | 5'-GCT TGG CGA TCT TCT TCT TG-3' |

### SUPPLEMENTARY METHODS

#### Fluorescence-activated cell sorting analysis

For surface marker staining, lymphocytes from ascites or blood were incubated with fluorochrome-conjugated antibodies against CD45, CD3, CD4, CD8, at the manufacturer's recommended dilution for 30 min at 4$^{o}$C as before[1]. To analyze intracellular markers like IFN-γ, IL4, Granzyme B and Perforin, lymphocytes were treated for 5 h with GolgiPlug (BD Biosciences, San Jose, CA) followed by surface staining with antibodies against CD4 or CD8. Cells were washed, fixed, and permeabilized with cytofix/cytoperm buffer (Proteintech, Rosemont, IL, USA) followed by incubation with various antibodies against intracellular markers. Flow cytometric analysis was performed on a BD FACS Calibur (BD Biosciences), and results were analyzed using Flowjo software (Version 10.8.1) (BD Biosciences). T-Distributed Stochastic Neighbor Embedding (tSNE) clusters were prepared using Flowjo software (Version 10.8.1)[2] as described before[2, 3]. All fluorochrome labeled antibodies (Table S2) were purchased from Biolegend (San Diego, CA).

#### Realtime PCR

Total RNA was extracted (Qiagen, Valencia, CA) from liver, tumor tissues, purified CD4+ and CD8+ T-cells and quantified by Qubit Fluorometer (Invitrogen, Waltham, MA). Reverse transcription was performed using 1 µg of total RNA using high-capacity cDNA kit in 20-µl reaction mixture and real-time polymerase chain reactions were performed and quantified as previously described[1] using CFX Bio-Rad Laboratories Real-time Polymerase Chain Reaction Detection system (Hercules, CA). Ribosomal protein L27 was used as a housekeeping gene. All primers were purchased from Integrated DNA Technologies (Coralville, IA). Primer sequence for ACAT, HMGCS2, HMGCL, BDH1 and GPR109a are listed in table 5.

#### Seahorse metabolic analysis

CD4$^{+}$ and CD8$^{+}$ cells were isolated from RD and IF tumor bearing mice using BD-IMAG anti- mouse CD4 and CD8 magnetic particles (BD Biosciences, San Jose, CA) and were plated at a density of 7 x10$^{5}$ cells/well in cell-tak coated XFe 96 cell plates. Oxygen consumption rate (OCR) and Extracellular acidification (ECAR) rate were measured using XFe 96 seahorse analyzer (Agilent Seahorse XF Analyzers) and analyzed as describe before[1, 4]. OCR measurements were recorded with port injections of (1) oligomycin (1 µmol), (2) FCCP (0.5 µmol), and a combination of (3) rotenone-antimycin at 1 µmol. ECAR, an indicator of aerobic glycolysis, was measured by incubating cells in an XFe base medium supplemented with 2 mmol/L glutamine. ECAR measurements were recorded after injecting with (1) glucose (10 mM), followed by (2) oligomycin (2 µM), and (3) 2-DG (100 mM). All media and coated plates were purchased from Agilent Technologies (Santa Clara, CA).

#### Quantitation of β-Hydroxybutyric acid (BHB) by LC-MS/MS

*Chemicals and reagents:* β-Hydroxybutyric acid standard was purchased from Sigma Aldrich (St Louis, MO) and isotopically labeled D-3-hydroxybutyrate ($^{13}$C4) used as an internal standard (ISTD), were purchased from Cambridge Isotope Laboratories (Tewksbury, MA). Acetonitrile, Water and Methanol and Formic acid, were purchased from Sigma Aldrich (St Louis, MO).

*Sample Preparation*: Concentrated stock solutions of β-Hydroxybutyric acid (β-HBA) standard (10 µg/mL) and the ISTD (250 µg/mL) were prepared in a 1:1 water: acetonitrile solution. Working solutions were prepared in matrix ranged from 15.75–1000 ng/mL for β-HBA. An ISTD working solution (0.500 ng/mL) was prepared in the extraction solvent 1:1 Water: acetonitrile. Calibration curve standards were prepared in duplicate for absolute concentration, while QC and blank (non-spiked) samples were prepared in Triplicates.

### Method validation

*Limit of detection (LOD) and Lower Limit of Quantification (LLOQ)*: Seven calibration standards ranging from (15.75,31.25,62.5,125,250,500 and 1000ng/ml) 15.75-1000ng/ml was subjected to the full extraction procedure three times before analysis. The limit of detection (LOD) was defined as the β-HBA concentration corresponding to the lowest calibration point, where signal to noise ratio (s/n) was three times greater than from the blank signal and lower limit of quantification was signal 10 times more compared to s/n with blank.

| Precursor | Selected MRM Precursor >Fragment | Collision energy | Cove voltage | Chromatographic Retention | |
|---|---|---|---|---|---|
| 105 | 105>87 | 15 | 10 | 1.10 | Primary |
| 105 | 105>43 | 15 | 8 | 1.10 | Confirmatory |
| 109 ($4C^{13}$) IS | 109>91 | 15 | 10 | 1.16 | Primary |

Selected β-HBA and its internal standard MRM transitions along with parameters and chromatographic retention.

| Time(min) | Flow Rate (mL/Min) | %A | %B | Curve |
|---|---|---|---|---|
| **0.0** | 0.3 | 90 | 10 | 6 |
| **2.0** | 0.3 | 60 | 40 | 6 |
| **3.0** | 0.3 | 40 | 60 | 6 |
| **3.5** | 0.3 | 40 | 60 | 6 |
| **3.8** | 0.3 | 90 | 10 | 6 |
| **5.0** | 0.3 | 90 | 10 | 6 |

Optimized gradient run for separation and quantitation of β-HBA in MRM mode. UPLC separation of β-HBA and its isotopically labeled internal standard (ISTD), using a Atlantis dC18, 130Å, 3 µm, 2.1 mm X 100 mm (P/N: 18601295)

*Data analysis:* Mass spectrometric data was acquired by MassLynx v4.2software. Quantification software: TargetLynxsoftware was used for preparing the calibration curve and absolute quantitation of β-HBA in the samples. Analyte concentrations were calculated using a 1/x weighted linear regression analysis of the standard curve.

*Quantitative Performance:* The quantitative performance using this sample preparation and LC-MS method was excellent, achieving a LLOQ of 15.75 ng/mL for β-HBA. Calibration curves were linear (r2 > 0.996) from 15.75–1000 ng/mL with accuracies between 85–115% with CVs.

*BHB extraction and LC-MS/MS analysis:* Experiment was setup for extraction and analysis conditions were optimized with RD, IF, control, β-BHA treatments. β-BHA absolute quantitation was performed in plasma and ascites. 100ul of plasma and ascites was used for extraction of the desired molecules using an in-house method for both the matrices. 300µl of acetonitrile was added to 100µl plasma and Ascites for protein precipitation and extraction of target molecule followed by 15 minutes of centrifugation at 15,000 rpm at $40^{0}$C. 350µl of upper layer was collated into fresh tube for complete drying under N2 based evaporation at room temperature. The dried residue was re-suspended in 500 µl diluent (Waters: Acetonitrile), vortexed and centrifuged (15 minutes at 15,000g at 4°C), and placed in an auto sampler vial for LC-MS/MS analysis. Waters UPLC-TQD mass spectrometry was employed for method development including the LC method optimization, ionization, and fragmentation tests. The binary pump was used to transport mobile phase A (Water+0.2% formic acid) and B (Acetonitrile+0.2% formic acid) at a flow rate of 0.3 ml/min in gradient mode. Best separation of β-Hydroxybutyric acid was achieved using UPLC with auto sampler with reversed-phase waters Atlantis dC18 Column, 130Å, 3 µm, 2.1 mm X 100 mm (P/N: 186001295) with in-line filter and guard kept at $50^{0}$C. A 5 minutes gradient employed an initial condition of 90% A with linear gradient to 60% A at 2.00 min, followed by linear gradient to 40% A at 3.50 min, followed by a return to 90% A in 0.3 minutes with 1.2 minutes re-equilibration with initial condition for β-Hydroxybutyric acid separation. The auto sampler was maintained at 4°C to compound stable and the injection volume was 5 µl with total running time of 5 minutes.

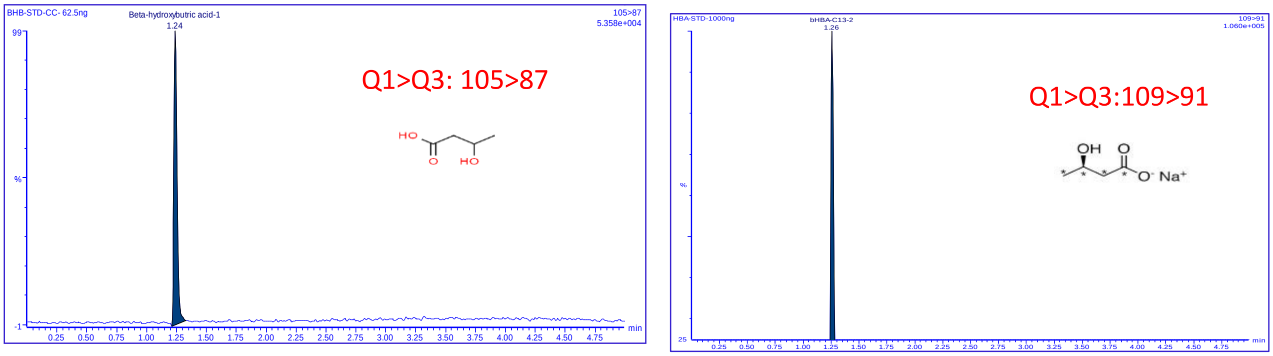


Figure 1: Typical MRM chromatogram of β-BHA (left panel) and IS (right panel) sample showing β-BHA peak.

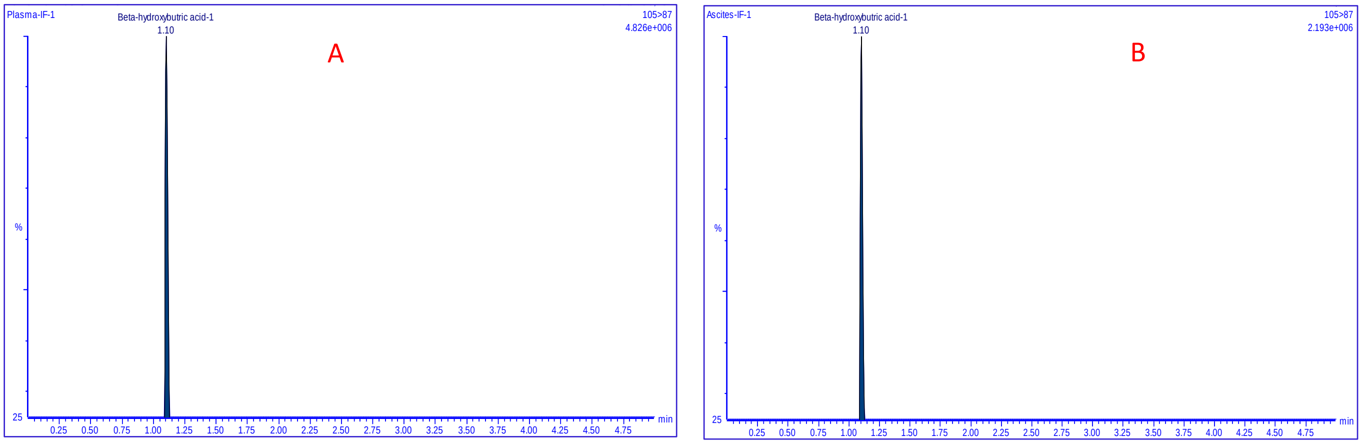


Figure 2: Panel A: Chromatography β-BHA (m/z 105>87) in plasma. Panel B: Extracted β-BHA in Ascites

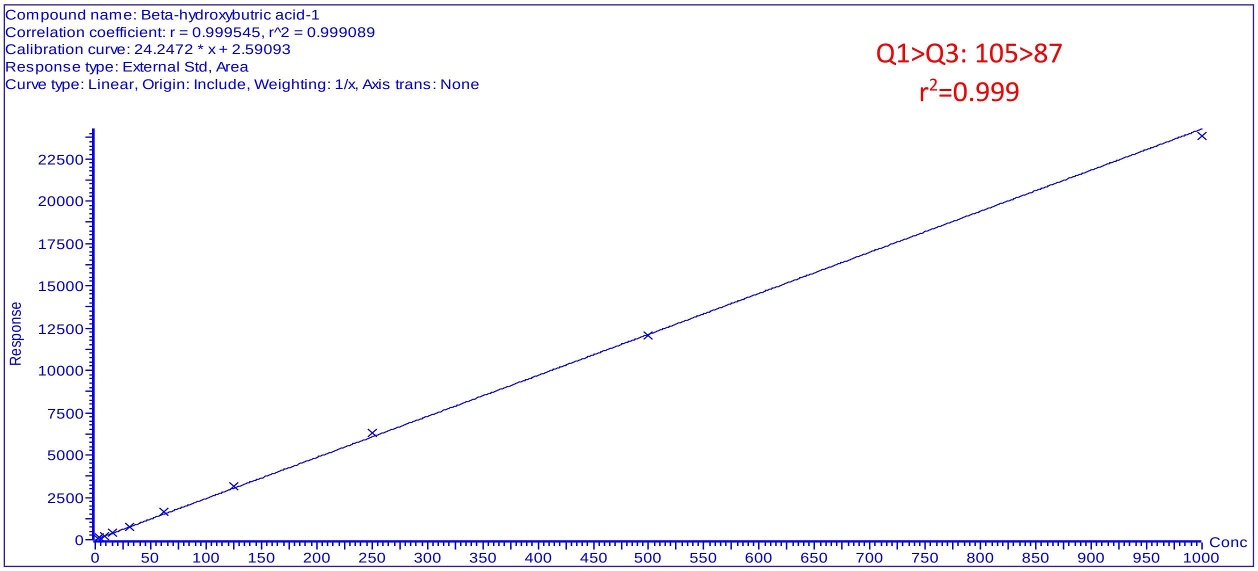


Figure 3: Seven point’s calibration curve ranging from (15.75-1000ng/ml) of β-BHA with r2=0.999

### SUPPLEMENTARY FIGURE LEGENDS

**Figure S1: IF does not reduce food intake or cause malnutrition.** Food intake every 24h was measured for a week, thrice during the study in mice eating ad libitum (RD) or eating during the 8h feeding window (IF) in mice injected with (A) ID8$^{p53+/+}$ or (B) ID8$^{p53-/-}$ or (C) ID8$^{p53-/-, PTEN-/-}$ tumors (n=6). Bar graph represents levels of albumin as a marker of malnourishment in plasma of RD and IF mice injected with (D) ID8$^{p53+/+}$ or (E) ID8$^{p53-/-}$ or (F) ID8$^{p53-/-, PTEN-/-}$ tumors, measured after 5 weeks of IF (n=5). NS= non-significant. IF compared to RD by Students T-test.

**Figure S2: IF decreases tumor promoting growth and proinflammatory factors in EOC mouse models.** Growth factors and proinflammatory cytokine levels were measured in ascites of RD and IF mice injected with ID8$^{p53-/-}$ cells. (A) Adipokine array from R& D biosystems was performed in ascites collected from RD and IF mice at 5 weeks and the top 6 significant proteins are highlighted with red rectangles. (B) Dot plot of significantly altered proteins ($p \leq 0.05$) in IF relative to RD. Ascites pooled from 4 mice per group was used to perform the array. (C) Bar graph represents fold-change of growth factors and inflammatory cytokines significantly altered in IF over RD ascites. (D) Table shows fold change of top 6 proteins observed in ascites of IF mice.

**Fig S3: IF reduces growth factors and proinflammatory factors in ID8$^{p53-/-, PTEN-/-}$ EOC model.** Growth factors and proinflammatory cytokines were measured in ascites (A-L) and plasma (M-X), collected at week 5 from ID8$^{p53-/-, PTEN}$ EOC bearing mice being subjected to RD or IF (n=4/group). (A, M) IGF-1, (B, N) Insulin, (C, O) Leptin, (D, P) Adiponectin, (E, Q) IFN γ, (F, R) IL-10, (G, S) TNFα, (H, T) IL-6, (I, U) IL-4, (J, V) IL-1β, (K, W) MCP-1 and (L, X) GMCSF levels measured by ELISA (n=4). *$p < 0.05$, **$p < 0.01$, ***$p < 0.001$, ****$p<0.0001$, assessed by Students t test.

**Fig S4: IF remodels the antitumor T cell responses in ID8$^{p53-/-, PTEN-/-}$ model.** Female C57/BL6 mice were injected with ID8$^{p53-/-, PTEN}$ cells intraperitoneally and subjected to RD and IF, ascites and were collected at 5 weeks from RD and IF mice were analyzed by flow cytometry (n=4). (A) A t-SNE visualization of markers after gating on single, live, CD45+ CD3+, CD4+, CD8+, IFNγ, IL4, Granzyme B (Gnzm b) and perforin expression. (B) Histograms represent average counts of the main T cell subsets in RD and IF groups. (C) Heat map represents marker expression of the main T cell subsets in individual samples. Bar graph represents the percentage of T cell subsets (D) CD4+, (E) CD4+IFNγ+, (F) CD+IL4+, (G) ratio of CD4+IFNγ+ to CD4+IL4+, (H) CD8+, (I) CD8+IFNγ+, (J) CD8+Gnz B and (K) CD8+perforin+. *$p < 0.05$, **$p < 0.01$, ***$p < 0.001$, ****$p < 0.0001$, IF compared to RD group, assessed by students t test.

**Fig S5: IF remodels the systemic anti-tumor T cell response:** Whole blood collected from RD and IF mice with ID8$^{p53-/-}$ or ID8$^{p53-/-, PTEN-/-}$ EOC were analyzed by flow cytometry (n=4). Bar graph represents the percentage of (A, I) CD4+, (B, J) CD4+IFNγ+, (C, K) CD+IL4+, (D, L) ratio of CD4+IFNγ+ to CD4+IL4+, (E, M) CD8+, (F, N) CD8+IFNγ+, (G, O) CD8+Grz B+ and (H, P) CD8+perforin+. *$p < 0.05$, **$p < 0.01$, ***$p < 0.001$, ****$p < 0.0001$, IF compared to RD group, assessed by Students t test.

**Fig S6: Anti-tumor effect of IF is T cell dependent:** ID8$^{p53-/-}$ cells were injected intraperitoneally into Nude mice (Nu). After 1 week, mice were subjected to RD or IF. (A) Levels of IGF-1, insulin, leptin, and adiponectin were measured in ascites by ELISA (n=4) (B) Cytokines were measured in ascites of RD and IF mice, including TNFα, IL-6, IL-4, IL-1β, MCP-1, GMCSF, IFNγ and IL-10 by ELISA (n=4) after 5 weeks. Female C57/BL6 mice were injected with ID8p$^{53-/-}$ cells intraperitoneally and subjected to RD, IF, IF with CD4+ depletion, IF with CD8+ depletion and IF with IgG2b control. Bar graph represents decrease in percentage of CD4+ and CD8+ cells in blood (C, D) and ascites (E, F) after respective 4 weeks of depletion. Female C57/BL6 mice were injected with ID8$^{p53-/-}$ cells intraperitoneally and subjected to RD, IF, anti-PD1 and IF combined with anti-PD1 treatments. Control RD and IF mice received IgG2b treatments. (G) Kaplan Meier graphs indicating overall survival (n=12) in all depletion and control groups. (n=8). (H) Bar graph represents average ascites volume in RD, IF, CD4+ Iso and CD4+ Dep. (I) Bar graph represents average ascites volume in RD, IF, CD8 Iso and CD8 Dep mice. Flow cytometry analysis was performed in blood collected from RD, IF, Anti-PD1 and IF combined with anti-PD1 mice after 5 weeks of treatments. (J) Histograms represent average counts of the main T cell subsets in all groups (K) Heat map represents marker expression of the T cell subsets in individual samples. Bar graph represents the percentage of (L) CD8+, (M) CD8+IFNγ+ and (N) CD8+Grz B+ cells. *p < 0.05, **p < 0.01, ***p < 0.001, ****p < 0.0001, using one-way ANOVA, followed by Sidak multiple comparison test.

**Fig S7: IF induces ketosis.** (A) Female C57/BL6 mice were injected with ID8$^{P53-/-}$ cells intraperitoneally and subjected to RD and IF. Plasma was isolated from whole blood from RD and IF mice and sent for untargeted metabolic profiling (n=6/ group). (B) Principal component analysis (PCA) on the metabolomic profiles displayed a clear separated clusters of metabolites in RD and IF groups. (C) Table representing the Over Representation Analysis on metabolomics data from RD and IF group.

**Fig S8: BHB inhibits ID8$^{p53-/-, PTEN-/-}$ EOC and induces anti-tumor CD8+ T cell response.** ID8$^{p53-/-, PTEN-/-}$ cells were injected intraperitoneally into C57/B6 female mice. After 1 week, mice were treated with either BHB (300mg/kg bd wt x 3 doses/week) or vehicle PBS (control mice, C). (A) Kaplan Meier graphs indicating overall survival (n=12), p=0.0001 by Gehan-Breslow-Wilcox test. Bar graph represents (B) average abdominal circumference, and (C) average ascites volume. Immune profiling was performed at 5 weeks (n=4). A t-SNE visualization of markers after gating on single, live, CD45+ CD3+, CD4+, CD8+, IFNγ+, IL4+, and Granzyme B + (GrzB) cells in ascites (D) and blood (E). Bar graph represents the percentage of T cell subsets CD4+, (F, L) CD4+IFNγ+, (G, M) CD4+IL4+, (H, N) CD8+, (I, O) CD8+IFNγ+ and (J, P) CD8+Grz B+ (K, Q) *p < 0.05, **p < 0.01, ***p < 0.001, ****p<0.0001, BHB compared to C, assessed by Student t test.

**Fig S9: BHB improves systemic anti-tumor responses in ID8$^{p53-/-}$ EOC.** Whole blood collected from RD and IF mice with ID8$^{p53-/-}$ EOC mice treated or untreated with BHB and were analyzed by flow cytometry (A) A t-SNE visualization of markers after gating on single, live, CD45+ CD3+, CD4+, CD8+, IFNγ+, IL4+ and Granzyme B + (GrzB) cells. (B) Histogram represents counts of the T cell subsets in ascites of C and BHB treated mice. (C) Heat map represents marker expression of the T cell subsets. Bar graph represents the percentage of T cell subsets (D) CD4+, (E) CD4+IFNγ+, (F) CD+IL4+, (G) CD8+, (H) CD8+IFNγ+ and (I) CD8+Gnz B+. (J) Splenic naïve CD4+ cells after activation were treated or untreated with 10mM BHB and co-cultured with 7AAD labeled ID8$^{p53-/-}$ cells were analyzed for percentage of live and dead ID8$^{p53-/-}$ and CD4 cells using flow. (K) Bar graph represents percentage of dead ID8$^{p53-/-}$ tumor cells. (L) Bar graph represents percentage of CD4+ cells. (M) IFNγ levels measured in supernatant collected from co-culture by ELISA. *p < 0.05, **p < 0.01, ***p < 0.001, ****p<0.0001, BHB compared to C, assessed by Students t tests.

**Fig S10: Anti-tumor immune response of IF and BHB.** Whole blood collected from control (C, untreated), IF or BHB treatments in ID8$^{p53-/-}$ EOC was analyzed by flow cytometry (A) A t-SNE visualization of markers after gating on single, live, CD45+ CD3+, CD4+, CD8+, IFNγ+, IL-4+ and

Granzyme B+ (GrzB) cells. (B) Histogram represents average counts of the T cell subsets measured. Bar graph represents the percentages of T cell subsets (C) CD4+, (D) CD4+IFNγ+, (E) CD+IL4+, (F) CD8+, (G) CD8+IFNγ+, and (H) CD8+Grz B+. $^{*}p < 0.05$, $^{**}p < 0.01$, $^{***}p < 0.001$, $^{****}p < 0.0001$, by one way ANOVA, followed by Sidak multiple comparison test.

**Figure S1**

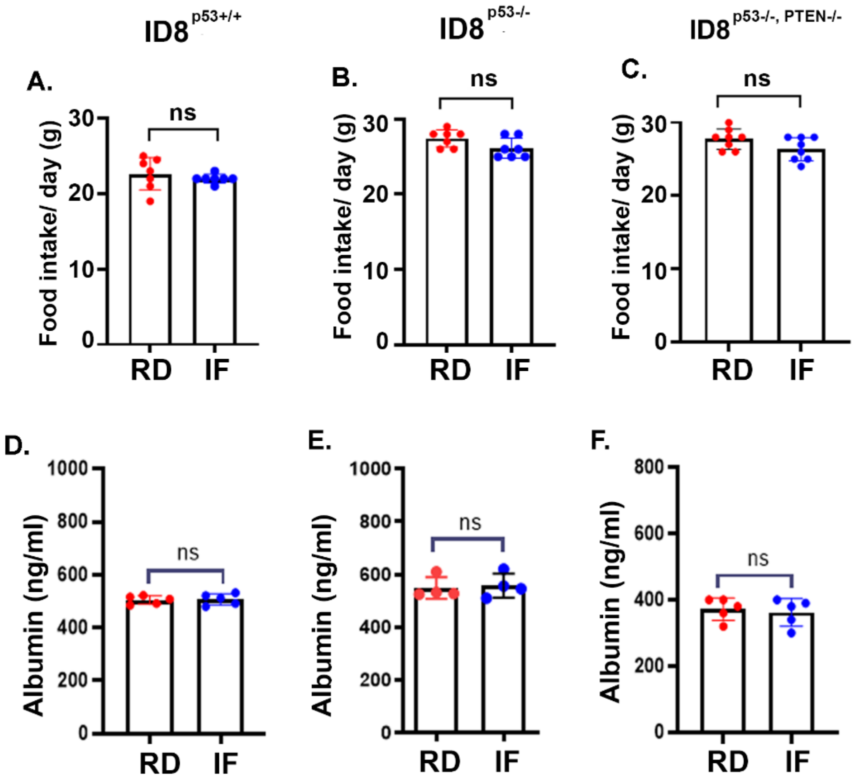

### Figure S2

**A.**

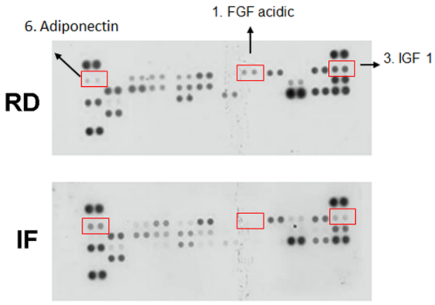


**B.**

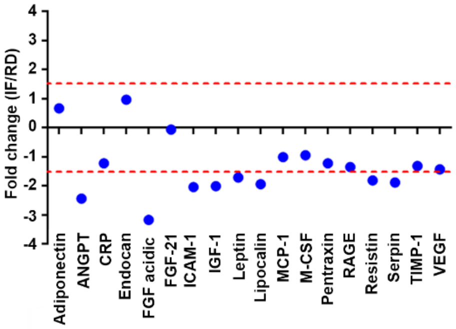


**C.**

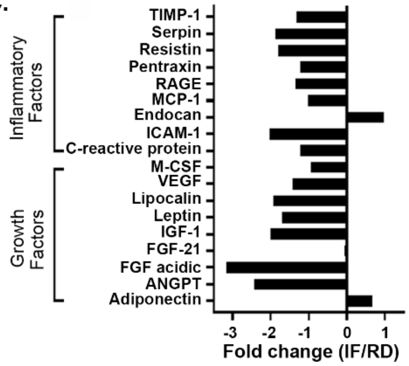


**D.**

| Top analyte | Fold change |
|---|---|
| 1. FGF acidic | -3.17 |
| 2. ANGPT | -2.44 |
| 3. IGF-1 | -2.01 |
| 4. ICAM 1 | -2.04 |
| 5. Lipocalin | -1.94 |
| 6. Adiponectin | +0.67 |

**Figure S3**

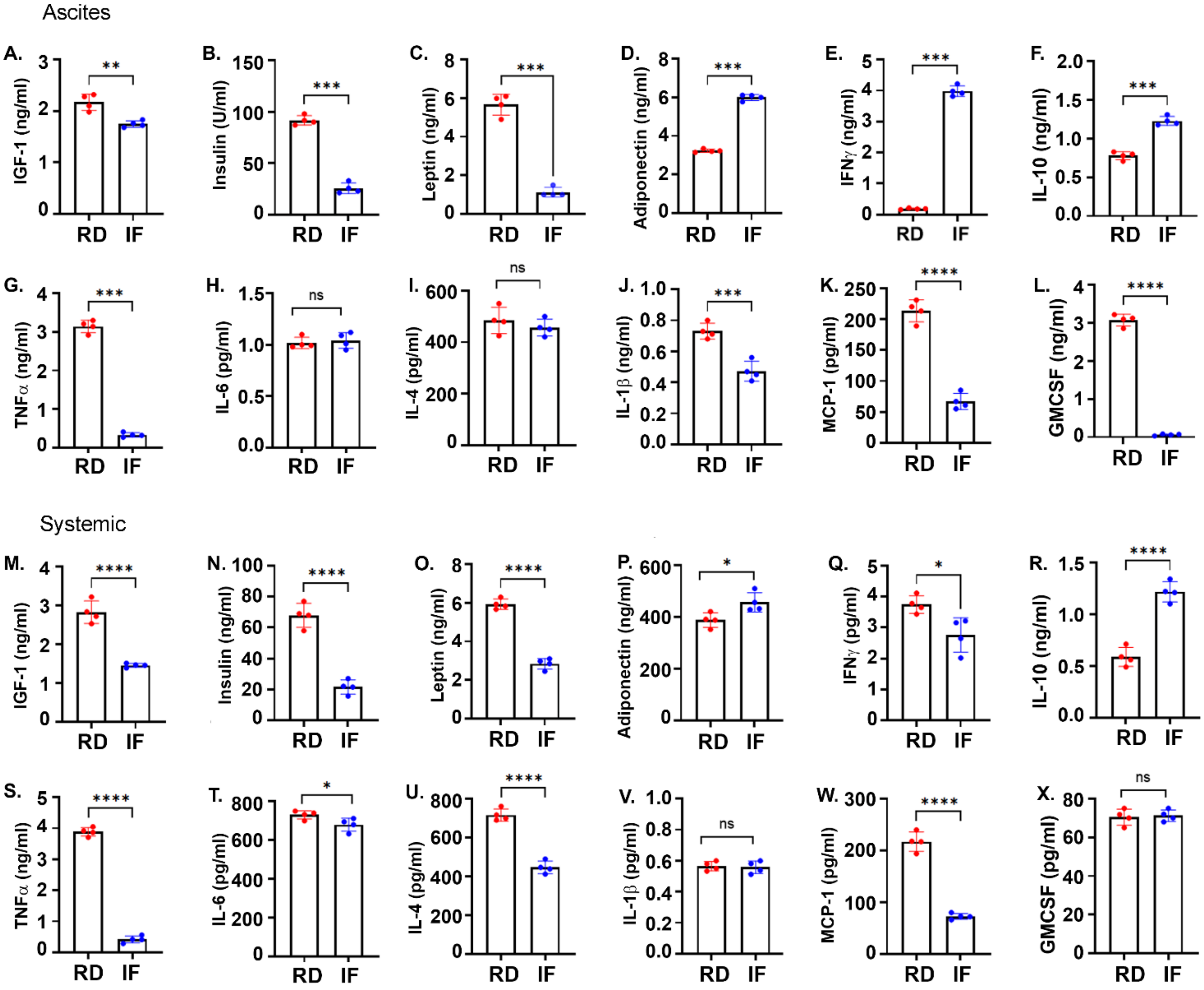
Ascites
A. IGF-1 (ng/ml) **
B. Insulin (U/ml) ***
C. Leptin (ng/ml) ***
D. Adiponectin (ng/ml) ***
E. IFNγ (ng/ml) ***
F. IL-10 (ng/ml) ***
G. TNFα (ng/ml) ***
H. IL-6 (pg/ml) ns
I. IL-4 (pg/ml) ns
J. IL-1β (ng/ml) ***
K. MCP-1 (pg/ml) ****
L. GMCSF (ng/ml) ****
Systemic
M. IGF-1 (ng/ml) ****
N. Insulin (ng/ml) ****
O. Leptin (ng/ml) ****
P. Adiponectin (ng/ml) *
Q. IFNγ (pg/ml) *
R. IL-10 (ng/ml) ****
S. TNFα (ng/ml) ****
T. IL-6 (pg/ml) *
U. IL-4 (pg/ml) ****
V. IL-1β (pg/ml) ns
W. MCP-1 (pg/ml) ****
X. GMCSF (pg/ml) ns
RD
IF

**Figure S4**

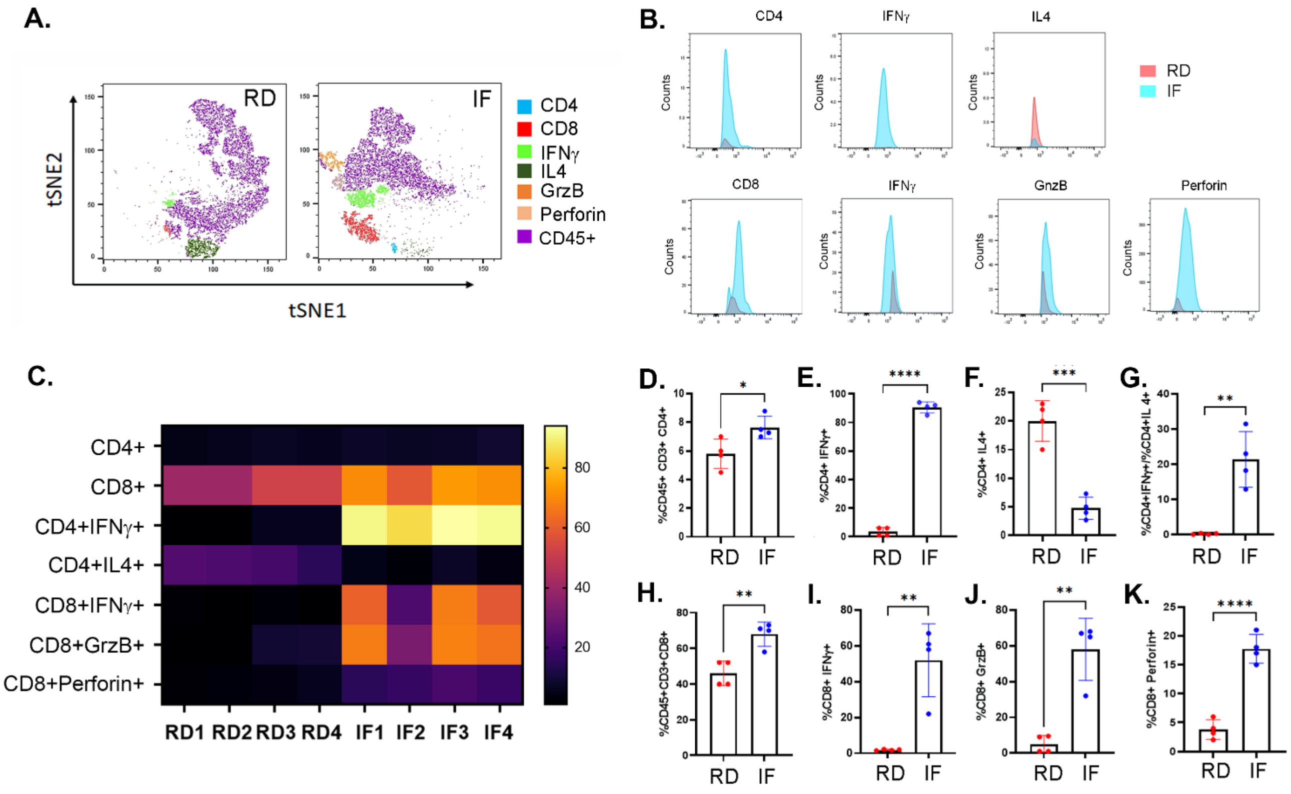
A.
RD
IF
tSNE2
tSNE1
CD4
CD8
IFNγ
IL4
GrzB
Perforin
CD45+
B.
CD4
IFNγ
IL4
RD
IF
CD8
IFNγ
GnzB
Perforin
Counts
C.
CD4+
CD8+
CD4+IFNγ+
CD4+IL4+
CD8+IFNγ+
CD8+GrzB+
CD8+Perforin+
RD1 RD2 RD3 RD4 IF1 IF2 IF3 IF4
80
60
40
20
D.
%CD45+ CD3+ CD4+
E.
%CD4+ IFNγ+
F.
%CD4+ IL4+
G.
%CD4+IFNγ+/%CD4+IL 4+
H.
%CD45+CD3+CD8+
I.
%CD8+ IFNγ+
J.
%CD8+ GrzB+
K.
%CD8+ Perforin+
RD
IF

**Figure S5**

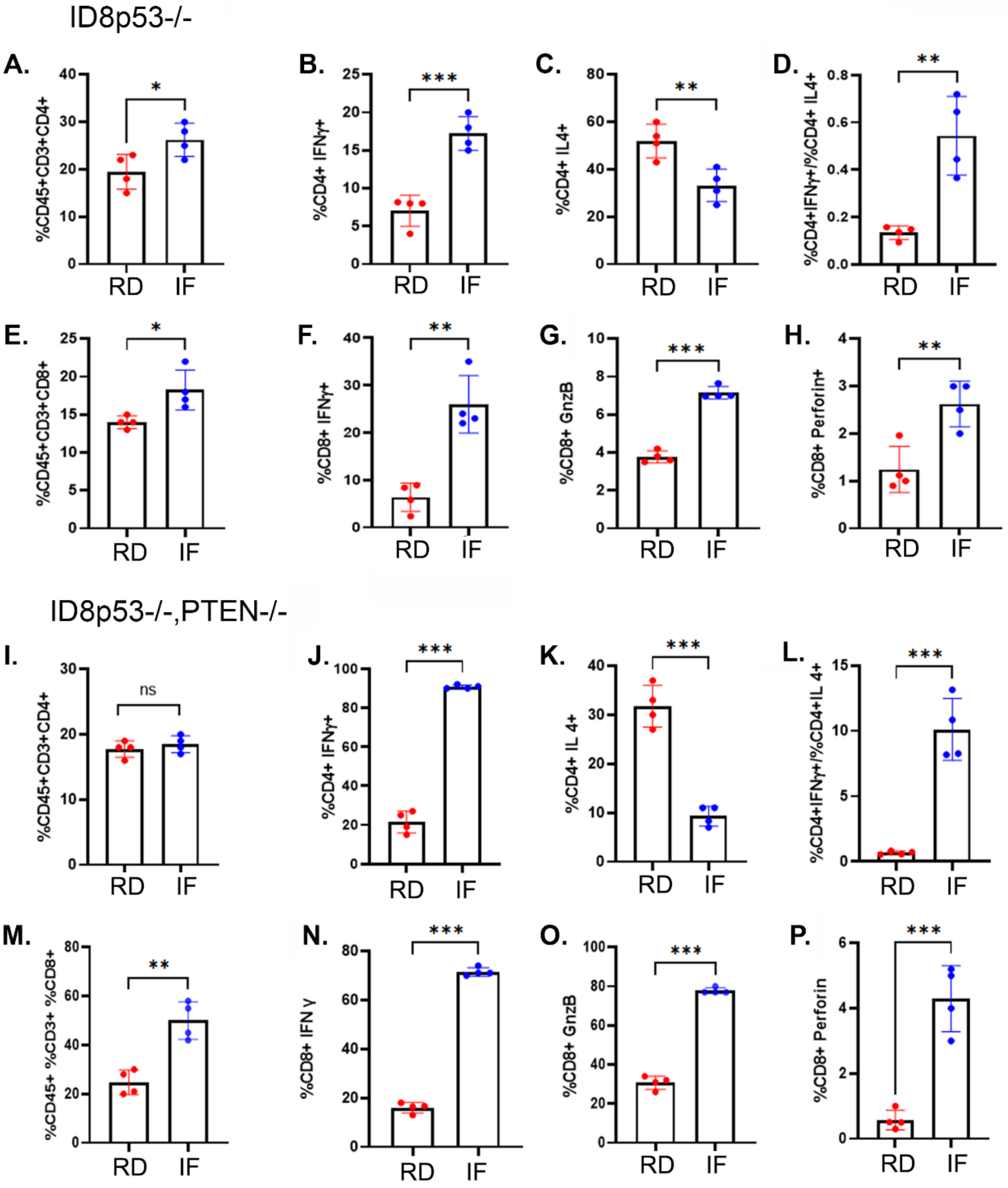
ID8p53-/-
A.
%CD45+CD3+CD4+
RD
IF
*
B.
%CD4+ IFNγ+
***
C.
%CD4+ IL4+
**
D.
%CD4+IFNγ+/%CD4+ IL4+
**
E.
%CD45+CD3+CD8+
*
F.
%CD8+ IFNγ+
**
G.
%CD8+ GnzB
***
H.
%CD8+ Perforin+
**
ID8p53-/-,PTEN-/-
I.
%CD45+CD3+CD4+
ns
J.
%CD4+ IFNγ+
***
K.
%CD4+ IL 4+
***
L.
%CD4+IFNγ+/%CD4+IL 4+
***
M.
%CD45+ %CD3+ %CD8+
**
N.
%CD8+ IFN γ
***
O.
%CD8+ GnzB
***
P.
%CD8+ Perforin
***

Figure S6

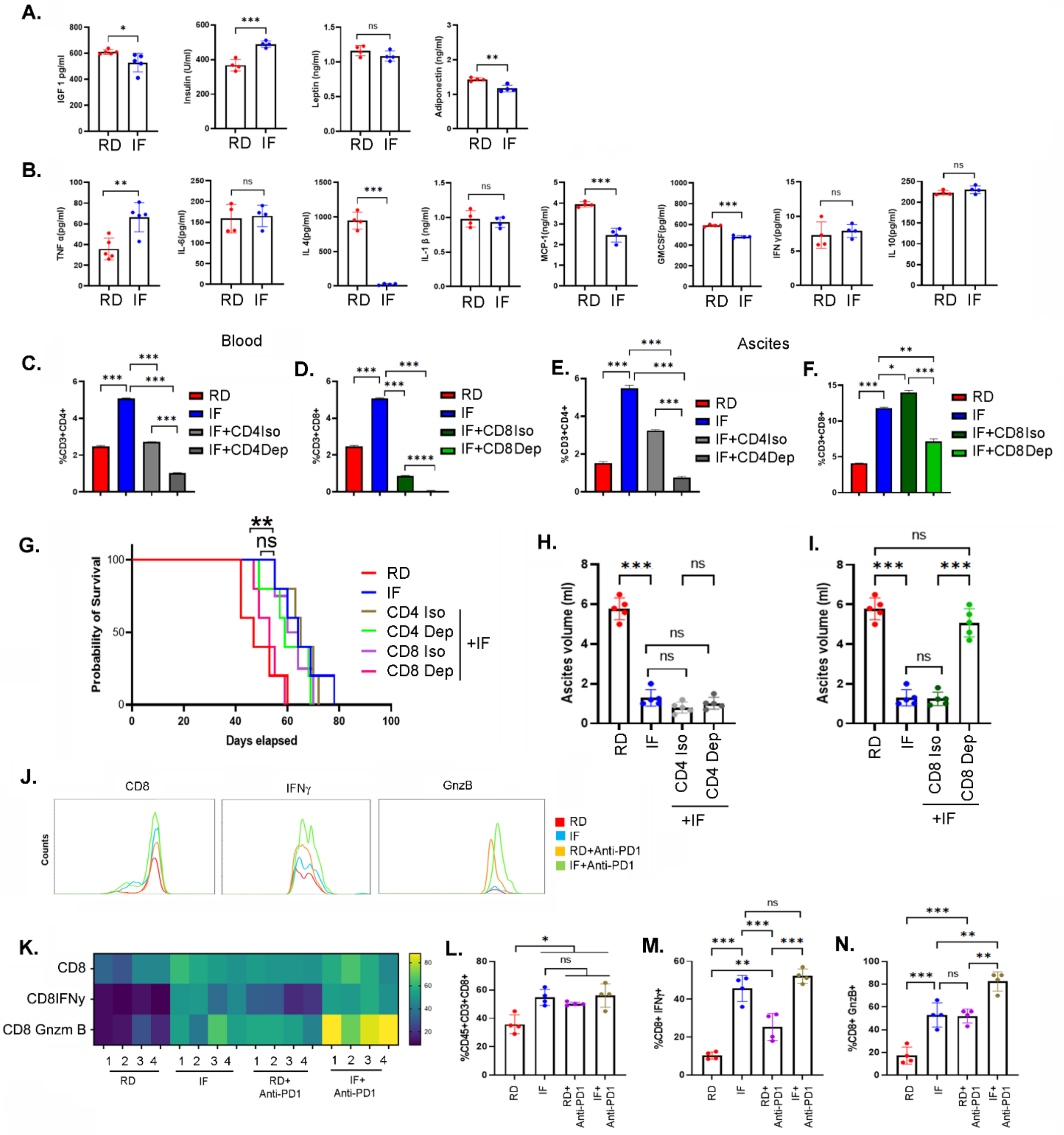

A.
IGF 1 pg/ml
Insulin (U/ml)
Leptin (ng/ml)
Adiponectin (ng/ml)
RD IF
B.
TNF α(pg/ml)
IL-6(pg/ml)
IL 4(pg/ml)
IL-1 β (ng/ml)
MCP-1(ng/ml)
GMCSF(pg/ml)
IFN γ(pg/ml)
IL 10(pg/ml)
Blood
Ascites
C.
%CD3+CD4+
RD
IF
IF+CD4Iso
IF+CD4Dep
D.
%CD3+CD8+
IF+CD8Iso
IF+CD8Dep
E.
F.
G.
Probability of Survival
Days elapsed
RD
IF
CD4 Iso
CD4 Dep
CD8 Iso
CD8 Dep
+IF
H.
Ascites volume (ml)
I.
J.
CD8
IFNγ
GnzB
Counts
RD+Anti-PD1
IF+Anti-PD1
K.
CD8
CD8IFNγ
CD8 Gnzm B
RD+ Anti-PD1
IF+ Anti-PD1
L.
%CD45+CD3+CD8+
M.
%CD8+ IFNγ+
N.
%CD8+ GnzB+

**Figure S7**

**A.**

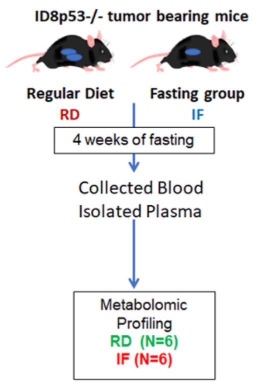


**B.**

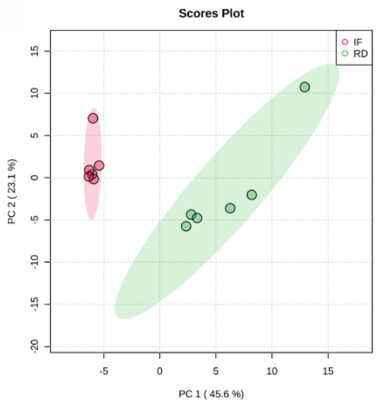


**C.**

Table 2: Result from Over Representation Analysis

| | total | expected | hits | Raw p | Holm p | FDR |
|---|---|---|---|---|---|---|
| Ketone Body Metabolism | 13 | 0.42 | 2 | 6.29E-02 | 1.00E+00 | 1.00E+00 |
| Fatty Acid Biosynthesis | 35 | 1.13 | 3 | 9.90E-02 | 1.00E+00 | 1.00E+00 |
| Beta Oxidation of Very Long Chain Fatty Acids | 17 | 0.55 | 1 | 4.30E-01 | 1.00E+00 | 1.00E+00 |
| Alpha Linolenic Acid and Linoleic Acid Metabolism | 19 | 0.61 | 1 | 4.66E-01 | 1.00E+00 | 1.00E+00 |
| Butyrate Metabolism | 19 | 0.61 | 1 | 4.66E-01 | 1.00E+00 | 1.00E+00 |
| Mitochondrial Beta-Oxidation of Medium Chain Saturated Fatty Acids | 27 | 0.87 | 1 | 5.92E-01 | 1.00E+00 | 1.00E+00 |
| Phenylalanine and Tyrosine Metabolism | 28 | 0.90 | 1 | 6.05E-01 | 1.00E+00 | 1.00E+00 |
| Methionine Metabolism | 43 | 1.39 | 1 | 7.63E-01 | 1.00E+00 | 1.00E+00 |
| Glutamate Metabolism | 49 | 1.58 | 1 | 8.07E-01 | 1.00E+00 | 1.00E+00 |
| Arginine and Proline Metabolism | 53 | 1.71 | 1 | 8.32E-01 | 1.00E+00 | 1.00E+00 |
| Pyrimidine Metabolism | 59 | 1.90 | 1 | 8.63E-01 | 1.00E+00 | 1.00E+00 |
| Valine, Leucine and Isoleucine Degradation | 60 | 1.93 | 1 | 8.68E-01 | 1.00E+00 | 1.00E+00 |
| Tyrosine Metabolism | 72 | 2.32 | 1 | 9.13E-01 | 1.00E+00 | 1.00E+00 |

**Figure S8**

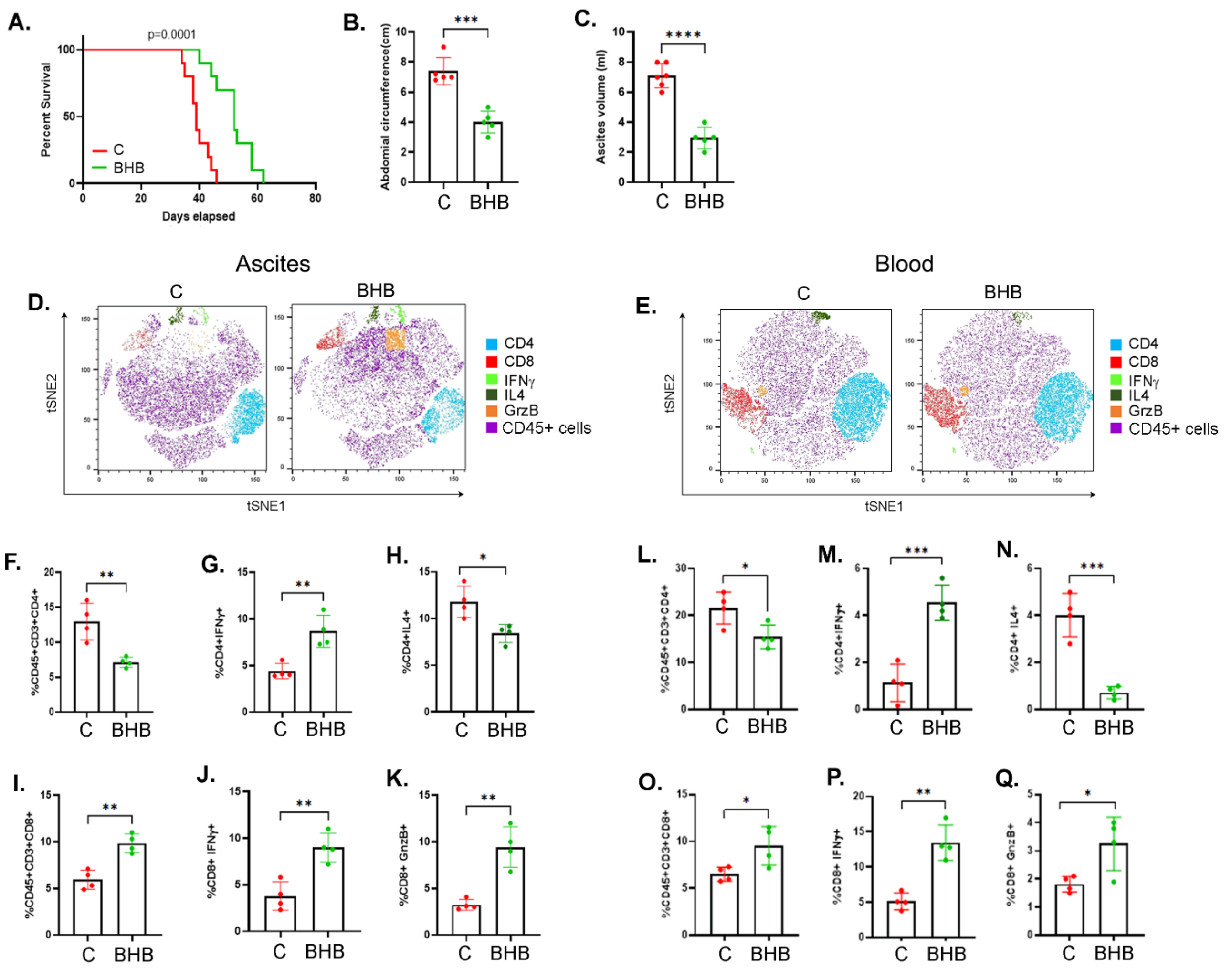

A.
p=0.0001
Percent Survival
Days elapsed
C
BHB
B.
Abdomial circumference(cm)
***
C.
Ascites volume (ml)
****
Ascites
Blood
D.
E.
tSNE2
tSNE1
CD4
CD8
IFNγ
IL4
GrzB
CD45+ cells
F.
G.
H.
L.
M.
N.
I.
J.
K.
O.
P.
Q.
%CD45+CD3+CD4+
%CD4+IFNγ+
%CD4+IL4+
%CD45+CD3+CD8+
%CD8+ IFNγ+
%CD8+ GnzB+

**Figure S9**

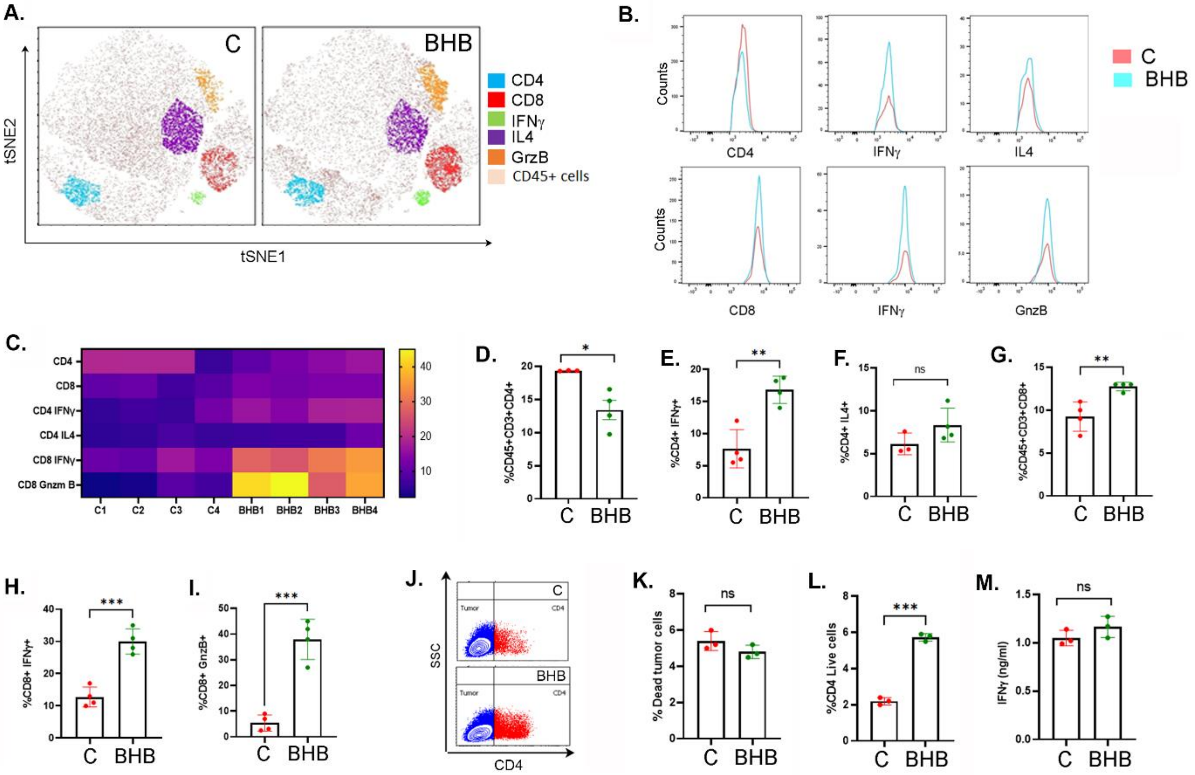

A.
C
BHB
CD4
CD8
IFNγ
IL4
GrzB
CD45+ cells
tSNE2
tSNE1
B.
C
BHB
Counts
CD4
IFNγ
IL4
CD8
GnzB
C.
CD4
CD8
CD4 IFNγ
CD4 IL4
CD8 IFNγ
CD8 Gnzm B
C1
C2
C3
C4
BHB1
BHB2
BHB3
BHB4
D.
%CD45+CD3+CD4+
E.
%CD4+ IFNγ+
F.
%CD4+ IL4+
G.
%CD45+CD3+CD8+
H.
%CD8+ IFNγ+
I.
%CD8+ GnzB+
J.
SSC
CD4
Tumor
K.
% Dead tumor cells
L.
%CD4 Live cells
M.
IFNγ (ng/ml)
ns

**Figure S10**

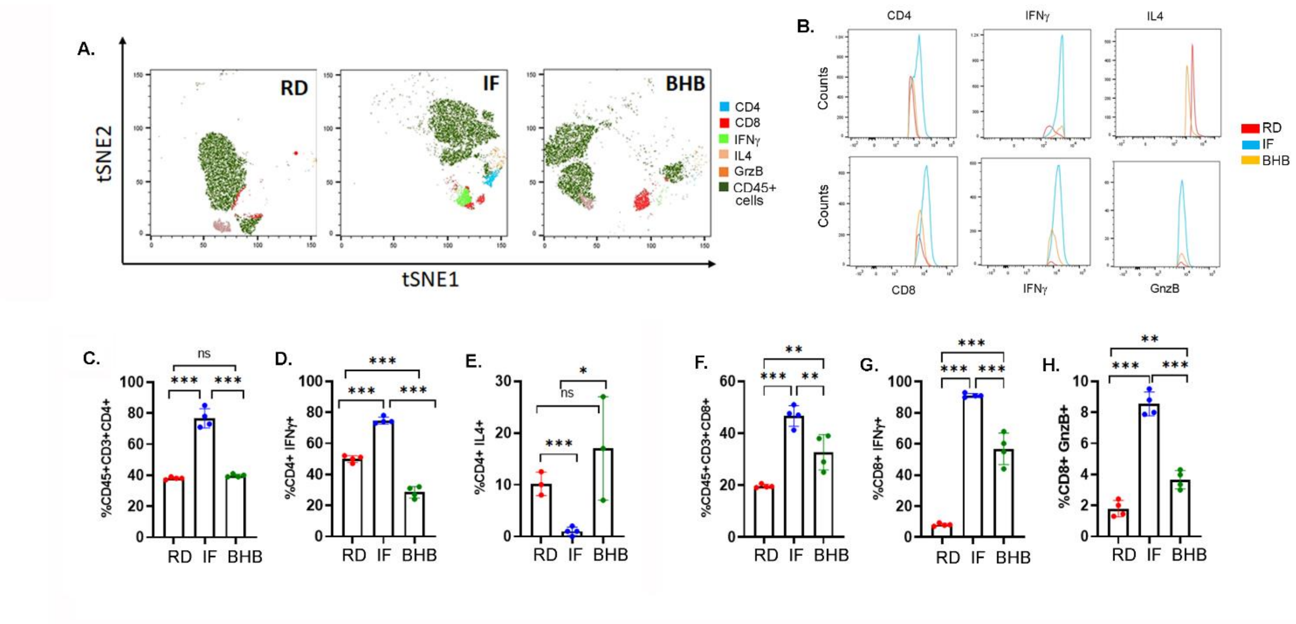
A.
RD
IF
BHB
tSNE2
tSNE1
CD4
CD8
IFNγ
IL4
GrzB
CD45+ cells
B.
CD4
IFNγ
IL4
Counts
CD8
GnzB
RD
IF
BHB
C.
%CD45+CD3+CD4+
D.
%CD4+ IFNγ+
E.
%CD4+ IL4+
F.
%CD45+CD3+CD8+
G.
%CD8+ IFNγ+
H.
%CD8+ GnzB+
ns
***
**
*
